# Dual inhibition of p38α MAPK and Casein kinase δ/ε as a novel treatment strategy for AR-independent and taxane-resistant advanced prostate cancer

**DOI:** 10.64898/2026.09.29.754128

**Authors:** Sayak Chakravarti, Suman Mazumder, Farnaz Hemmati, Farshad Amiri, Sarah Batten, Panagiotis Mistriotis, Robert Rusty Arnold, Ujjal Kumar Mukherjee, Taraswi Mitra Ghosh, Amit Kumar Mitra

## Abstract

Prostate cancer (PCa) is the most commonly diagnosed cancer and the second leading cause of cancer death among US men. Metastatic castration-resistant PCa (mCRPC) is a clinically advanced form of PCa, often associated with increased aggressiveness, metastatic potential, morbidity, and a higher risk of developing resistance to taxanes (TX), the first-line chemotherapy for mCRPC. Furthermore, cancer ‘stemness’ and epithelial to mesenchymal trans-differentiation (EMT) potentially contribute to aggressiveness and the development of drug resistance in mCRPC. Here, we applied our drug development pipeline, secDrug, which utilizes a pharmacogenomics data-driven computational algorithm, to demonstrate that TAK-715 - a dual inhibitor of p38α MAPK and Casein kinase δ/ε, is a promising treatment for these advanced/lethal variants of PCa.

Using *in vitro* cytotoxicity assays followed by cell-based functional assays in a panel of mCRPC cell lines representing TX-sensitive mCRPC, clonally derived TX-resistant lines, and a highly aggressive metastatic variant of mCRPC, we demonstrated the efficacy of TAK-715 treatment as a single agent and in combination with TX, including cancer ‘stem-like’ cells. Further, we showed that the apoptotic effects of TAK-715 occur via a mitochondrial-mediated pathway. Bulk tumor RNA sequencing followed by pathway analysis identified genes associated with mitochondrial dysfunction and cell cycle arrest as the top molecular networks associated with TAK-715 single-treatment. HES1, a gene associated with nodal metastasis and PCa progression, was the top differentially expressed gene. Single-cell transcriptomics analysis revealed that TAK-715 treatment erodes the subclonal populations responsible for cancer stemness, metastasis, and drug resistance. The clinical significance of these findings was validated *in silico* using multiple patient datasets.

Our results suggest that TAK-715 treatment has the potential to decrease oncogenic progression and cancer stem cell-like activity in drug-resistant, aggressive, and stem-like mCRPC cells.

**TEASER:** To address the unmet need for novel strategies to treat aggressive/resistant/metastatic (advanced) prostate cancer, this study demonstrates the efficacy of dual inhibition of p38α MAPK and Casein kinase δ/ε through a novel approach that integrates *in silico* analysis with RNA-seq, scRNA-seq, cell-based functional assays, and *ex vivo* validation.

## INTRODUCTION

Prostate cancer is the most common cancer in men in the USA, with ∼313,780 new cases (15.4% of all new cancer cases) and 35,770 deaths (5.8% of all cancer deaths) in 2025 (NCI SEER Cancer Stat Facts: Prostate Cancer, https://seer.cancer.gov/statfacts/html/prost.html)^1^. A critical requirement for PCa is the initial reliance on androgen receptor (AR) signaling for growth and to avoid apoptosis^2^. Androgen deprivation therapy (ADT or medical castration), achieved through surgical or pharmacological approaches, exploits this dependence of PCa on androgens and serves as the standard treatment option for early-stage PCa^3–5^. While most early-stage PCa patients (castration sensitive or CSPC) treated with ADT show good initial response, vast majority of these men eventually become unresponsive towards hormone therapy and, despite low levels of androgen, the disease progresses with continuously rising Prostate Serum Antigen (PSA), eventually developing more lethal/advanced forms called Castration-resistant prostate cancer (CRPC)^5–7^. Further, when CRPC becomes metastatic (mCRPC), it no longer remains localized in the prostate and spreads to other tissues and organs like bone, lymph nodes, bladder, liver, lung, and brain^8^. Metastatic castration-resistant prostate cancer (mCRPC) is the clinically most advanced and lethal disease state, with a 5-year median survival rate of 31%^9^. Although next-generation AR targeting chemotherapy and Taxane (TX)-based chemotherapy (Docetaxel (DTX)) confer modest overall survival benefits in chemotherapy-naïve mCRPC patients, eventual development of acquired drug resistance over time is nearly universal, where progression-free survival (PFS) approaches almost 0% in 3 years^10^. Moreover, variable drug response observed among patients (only ≈50% of men show measurable DTX response) due to innate or refractory resistance (resistance that is already present in drug-naive patients) is also a cause of concern in mCRPC chemotherapy^11^. Treatment with the second-generation anti-mitotic Taxane, Cabazitaxel (CBZ), in patients who have received prior DTX confers little improvement (≈2.4 months) in the median survival and causes severe side effects^12^. Chemotherapy options become limited once patients fail TX therapy. Further, ‘cancer stemness’ characterized by the presence of cancer stem-like cells (CSCs) like side populations and CD44+ cells, and the acquisition of mesenchymal phenotype or epithelial to mesenchymal trans-differentiation (EMT) with self-renewal and differentiation capacities, is believed to significantly contribute to the development of drug resistance^13^. Thus, drug development for mCRPC treatment poses a significant challenge with very few therapeutic successes. To serve this purpose, we have designed a novel pharmacogenomics data-driven optimization-regularization-based computational algorithm called “secDrug” to predict targets for novel secondary drugs (“secDrugs”) for the management of advanced cancers, including metastatic castration-resistant and taxane-resistant PCa subtypes^14,15^. We hypothesize that our predicted secDrugs for PCa will be useful in curbing oncogenic progressions and abrogate drug resistance and cancer ‘stemness’ through simultaneous inhibition of multiple oncogenic factors/pathways.

In this study, we demonstrate that the secDrug, TAK-715 - a small molecule inhibitor of p38 MAPK specific for p38α, as well as casein kinase δ/ε, is indeed effective against androgen-independent and TX-resistant mCRPC, as well as subclones representing ‘cancer stemness’. Further, we used bulk tumor RNA sequencing, single-cell transcriptomics, and functional validation to identify potential molecular networks/pathways associated with TAK-715 treatment and TAK-715+TX drug synergy, which were validated *in silico* across multiple patient datasets.

## RESULTS

### TAK-715 as the top secDrug for mCRPC therapy

We have designed the secDrug algorithm, a pharmacogenomics data-driven *in silico* drug prediction pipeline that applies a greedy algorithm-based set-covering computational optimization method, followed by a regularization technique to predict novel targets for secondary drug candidates (“secDrugs”) against drug-resistant and advanced cancers^14^. When applied to genitourinary cancers, including PCa, our secDrug algorithm, we identified the following drug targets for small molecular inhibitors that could be used as secondary drugs to kill the maximum number of advanced PCa cell lines when used in combination with the primary drug, Taxane: NAMPT, p38α MAPK, BIRC5, Akt1/Akt2/Akt3, PKCβ, Wnt/β-catenin, MEK1/2, and EGFR ^14^. Among these, we have shown earlier that the NAMPT inhibitor, FK866, is indeed effective against advanced, taxane-resistant, and stem-like cells in lethal PCa^14^.

### TAK-715 as a dual inhibitor of p38α MAPK and Casein kinase δ/ε

TAK-715 is a small-molecule inhibitor of p38α MAP Kinase. Using a fluorescence-based Z-Lyte Kinase assay, we found that TAK-715 inhibits multiple kinases in addition to p38α MAPK (MAPK14). **Figure 1A** shows that TAK-715 also targets MAP4K4 (hepatocyte progenitor kinase-like/germinal center kinase-like kinase (HGK) and Nck-interacting kinase (NIK) and the Casein Kinases δ (CSNK1D) and ε (CSNK1E).

**Figure 1A.**
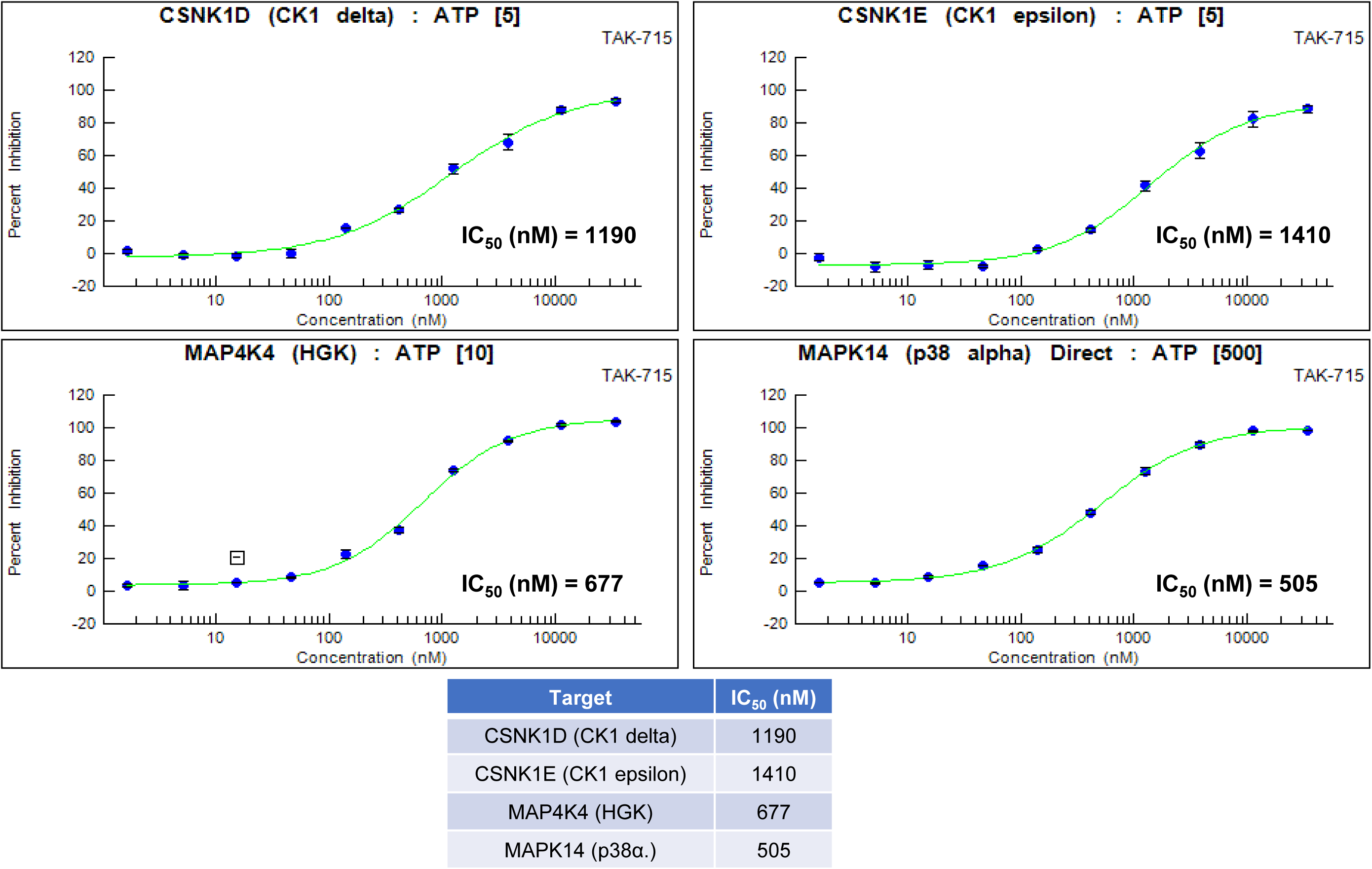
TAK-715 inhibits p38α MAPK and Casein kinase δ/ε. Z’LYTE Kinase Assay data showing kinetic study of TAK-715 with varied concentrations to determine IC_50_ against different target kinases.

The TAK-715 target genes were also confirmed using the Harvard Medical School (HMS) ‘s NIH Library of Integrated Network-based Cellular Signatures (LINCS) perturbagen database, a publicly available database devoted to understanding how human cells respond to perturbation by drugs, the environment, and mutation^16^.

Next, we used single-cell RNA sequencing (scRNA-seq) as a biomarker-based secDrug screening tool to identify single-cell sub-clones in mCRPC cell lines harboring TAK-715 target genes. We have earlier observed that our scRNA-seq data on the untreated cell lines showed that the majority of t-distributed stochastic neighbor embedding (t-SNE) clusters/subclones have enrichment of genes that play a major role in cancer progression, development, and maintenance of cancer stemness^14^. Interestingly, these subclusters also showed high expression of the target genes MAPK14, MAP4K4, CSNK1D, and CSNK1E (**Figure 1B**), indicating that TAK-715 may be effective against these taxane-resistant, stem-cell-like subclusters.

**Figure 1B.**
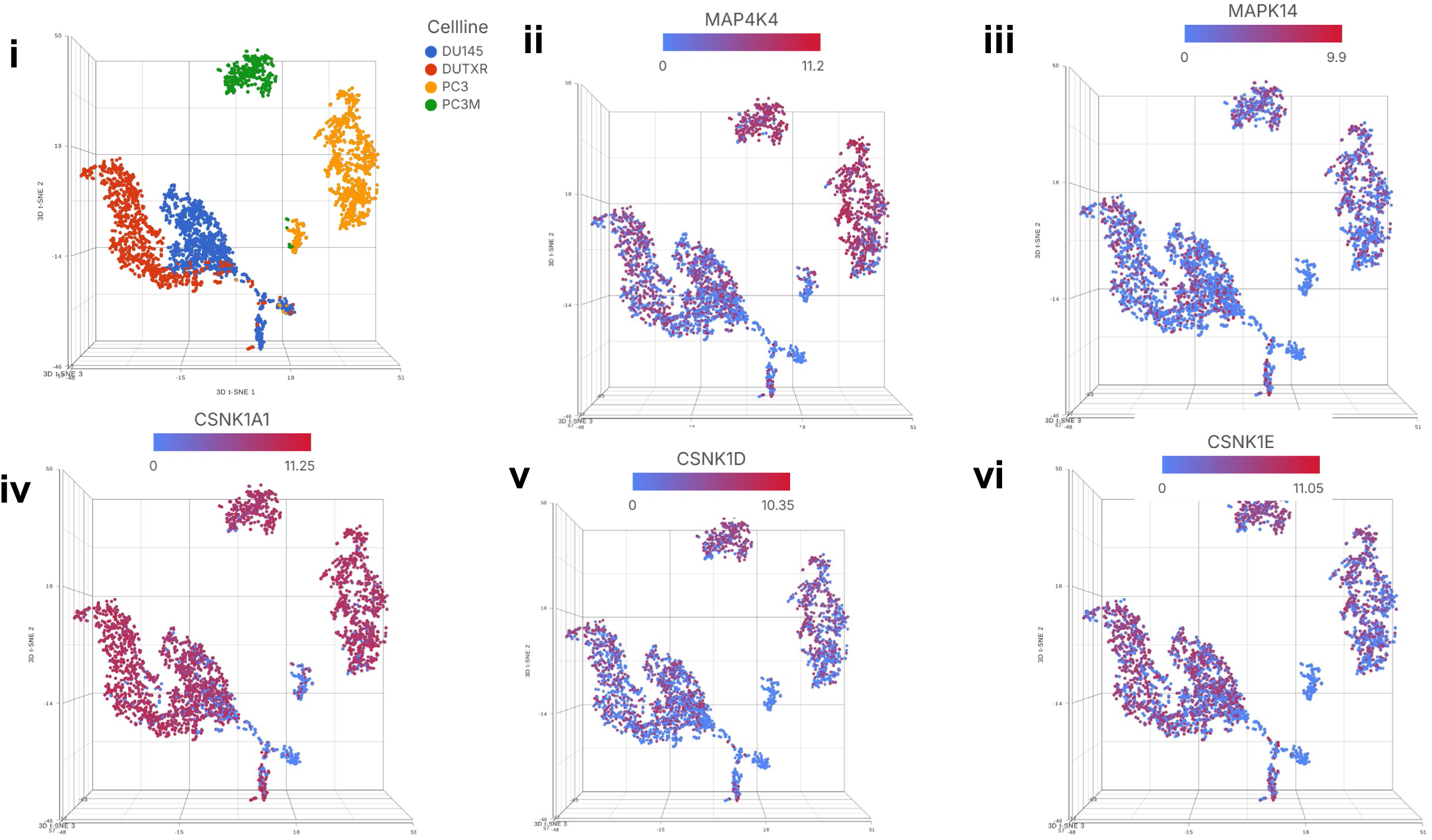
Single cell transcriptomics of human mCRPC cell lines reveals TAK-715 is effective against mCRPC, TX-resistant and stem-cell like sub-clones. **i.** tSNE plot showing untreated and TAK7-715-treated DU145 cells. **ii-vi.** PCa subclusters showed high expression of potential TAK-715 target genes. TAK-715 target genes were derived from Z’-Lyte assay (Figure 1) and the Harvard Medical School (HMS)’s NIH Library of Integrated Network-based Cellular Signatures (LINCS) perturbagen database. Single-cell RNA sequencing using the Droplet sequencing method (10X Genomics) was performed on the PCa cell lines DU145, DUTXR, PC3, PC3M. t-distributed stochastic neighbor embedding (t-SNE) plots showing the comparison between the single-cell clusters.

### TAK-715 induces reduction of cellular viability as a single agent and shows synergy with standard-of-care PCa drugs

We determined the cytotoxic effects of TAK-715 on a panel of the AR-ve mCRPC cell lines represented by DU145, PC3, the clonally-derived acquired taxane-resistant lines DUTXR and PC3-TXR (>50 fold higher TX IC_50_ compared to parental lines), and PC3M – the more aggressive and metastatic subline of PC3. Single-agent survival curves showed that TAK-715 was effective against all mCRPC cell lines and significantly reduced the number of viable cells in a dose-dependent manner (**FIGURE S1**). Furthermore, single-agent IC_50_ values for DTX and CBZ were negatively correlated with the IC_50_ value of TAK-715 in these cell lines (p<0.05). The correlation coefficients are -0.2 for DTX and TAK-715, and -0.4 for CBZ and TAK-715.

Next, we investigated the impact of using combination therapy regimens consisting of a constant dose of TAK-715 as well as 3 different doses equivalent to the IC_50_, IC_25_, and IC_50_/2 values of TAK-715 in each cell line and a range of DTX concentrations (0.2-400 nM for DU145, PC3 & PC3M; 50-2250 nM for DUTXR and PC3-TXR). All combinations showed a significant reduction in cellular proliferation compared with DTX alone, indicating drug synergy (**Figure 2A**). Evidently, the combination index (CI) values, a measure of the synergistic action between DTX and TAK-715 in combination, were consistently <1, indicating very high synergy. Similar results were observed for the CBZ and TAK-715 combination treatment (**Figure 2B**). The synergistic effects were particularly profound in DUTXR, PC3-TXR, and PC3M.

**Figure 2A.**
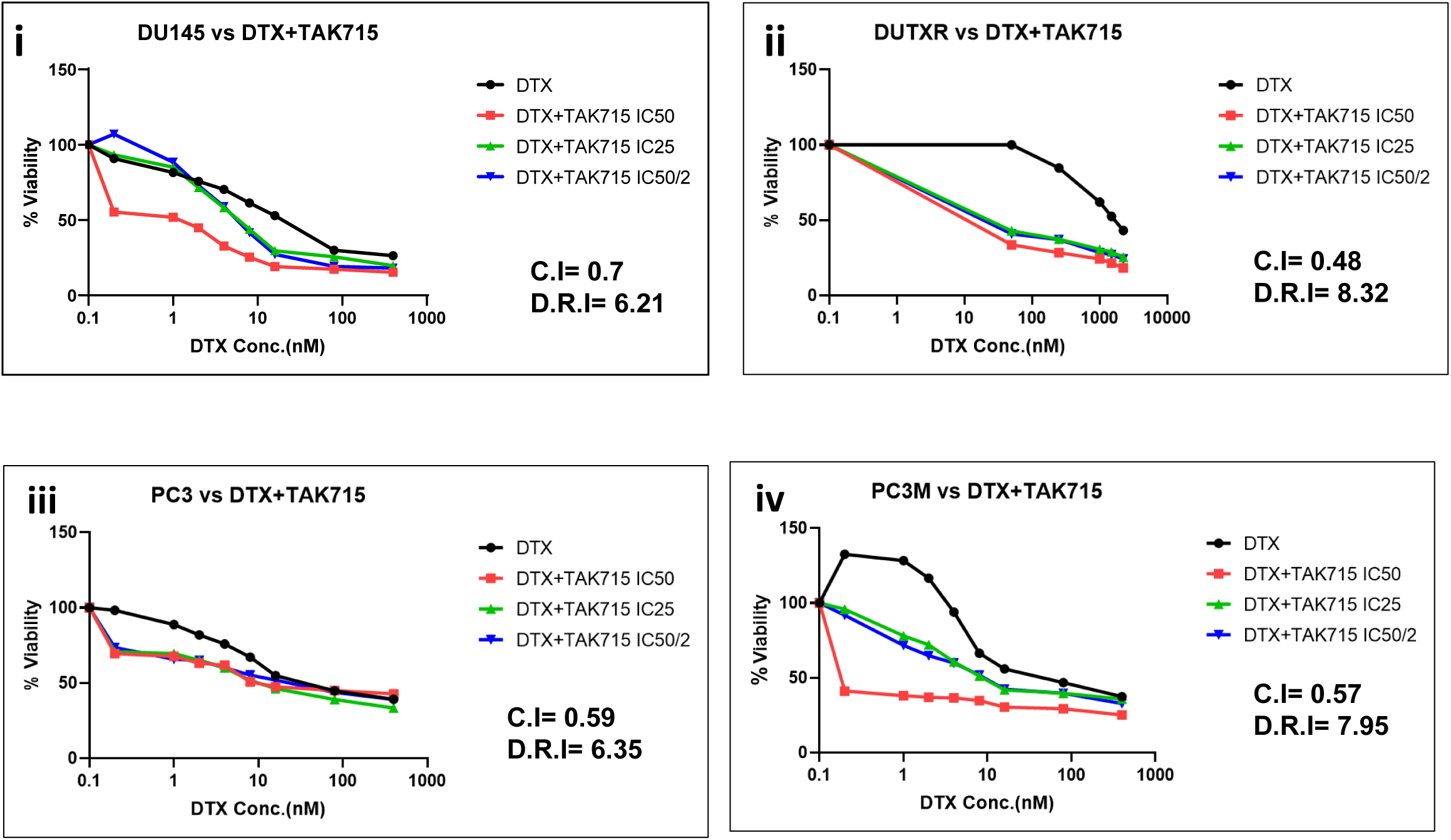
Docetaxel + TAK-715 combination therapy exhibits synergy in mCRPC. *In vitro* cell viability profile of AR-ve mCRPC cell lines (i) DU145, (ii) DUTXR, (iii) PC3 (iv) PC3M treated, with treated with different combination of Docetaxel+TAK-715.

**Figure 2B.**
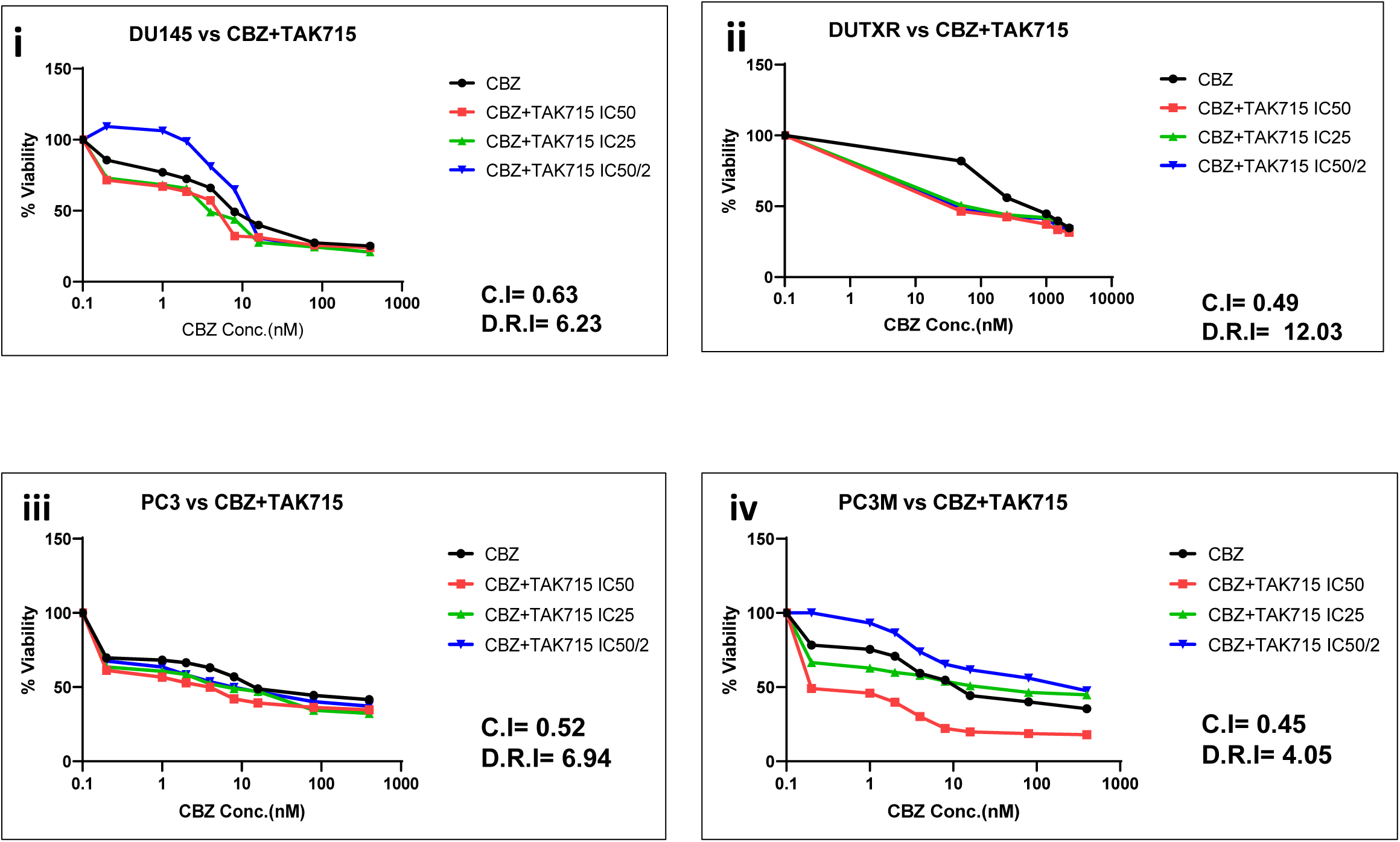
Cabazitaxel + TAK-715 combination therapy exhibits synergy in mCRPC. *In vitro* cell viability profile of AR-ve mCRPC cell lines (i) DU145, (ii) DUTXR, (iii) PC3 (iv) PC3M treated with treated with different combination doses of Cabazitaxel+TAK-715.

To further validate the synergism of TAK-715+DTX, we performed live-cell imaging to monitor changes in morphology and live-cell count. Cells were treated with DTX and TAK-715 alone or in combination with the two drugs for 48 hours. We observed that combining IC25 doses of both drugs reduced the average live cell count by 76±9% relative to the control and by 52±17% relative to DTX single-agent treatment (**FIGURE S2A**). Further, the average reduction in live cell count for the combination treatment with IC_50_ doses of both drugs was 86±4% compared to the control and 73±8% compared to the IC_50_ dose of DTX as single-agent treatment. Dose-dependent cell shrinkage, or a decrease in cell volume, was also observed, a ubiquitous feature of programmed cell death.

Further, an assessment of the nuclear morphology of attached cells using NucBlue staining, a reagent commonly used to distinguish condensed nuclei in apoptotic cells, revealed TAK-715-induced morphological changes, such as nuclear fragmentation and chromatin condensation, indicative of apoptosis **(Figure S2B)**. In addition, our results showed even higher cell death and more nuclear damage in the TAK-715+DTX combination treatment compared to the TAK-715 single-drug treatment.

### DTX+TAK-715 combination enhances apoptosis in AR-ve mCRPC

The impact of TAK-715 on cellular apoptosis was assessed using Caspase 3/7 Glo Assay. We observed significantly elevated Caspase 3/7 activity following treatment with TAK-715 and primary PCa drugs as single agents, indicating increased apoptosis, which was more pronounced when the drugs were used in combination than with individual single-agent treatments (**FIGURE S3A-B**).

### TAK-715 potentially reduces cell migration and tumor metastasis

Next, we performed a novel microfluidic chamber-based cell migration assay, coupled with time-lapse microscopy, to assess the effect of our drug combination on the migratory potential of AR-ve mCRPC cells. This microfluidic chamber represents a physiologically relevant model for the *in vitro* study of cell motility through confining microenvironments of different dimensions^17^. Our results showed that TAK-715 single-agent and DTX + TAK-715 combination treatment reduced cell entry into moderately confining channels, indicating a generic defect in the mechanisms of cell migration, PCa cell invasion, and potentially tumor metastasis (loss of plasticity and metastatic potential) (**FIGURE 3A**). To validate the effect of the TAK-715+DTX combination on cellular motility and migration, a wound healing assay was performed. A scratch was made in the cell monolayer, and the cells were treated with TAK-715 as single agents and in combination with DTX. Image analysis of the scratches using the ImageJ software showed an average reduction of 60±13% in the wound area in the cells of the control well after 24 hours. In contrast, there was only 8.5±2.5% reduction in the drug combination-treated cells. After 48 hours, it was 81±2% and 18±8%, respectively (**FIGURE 3B**), signifying a higher effect in reducing cell migration.

**Figure 3A:**
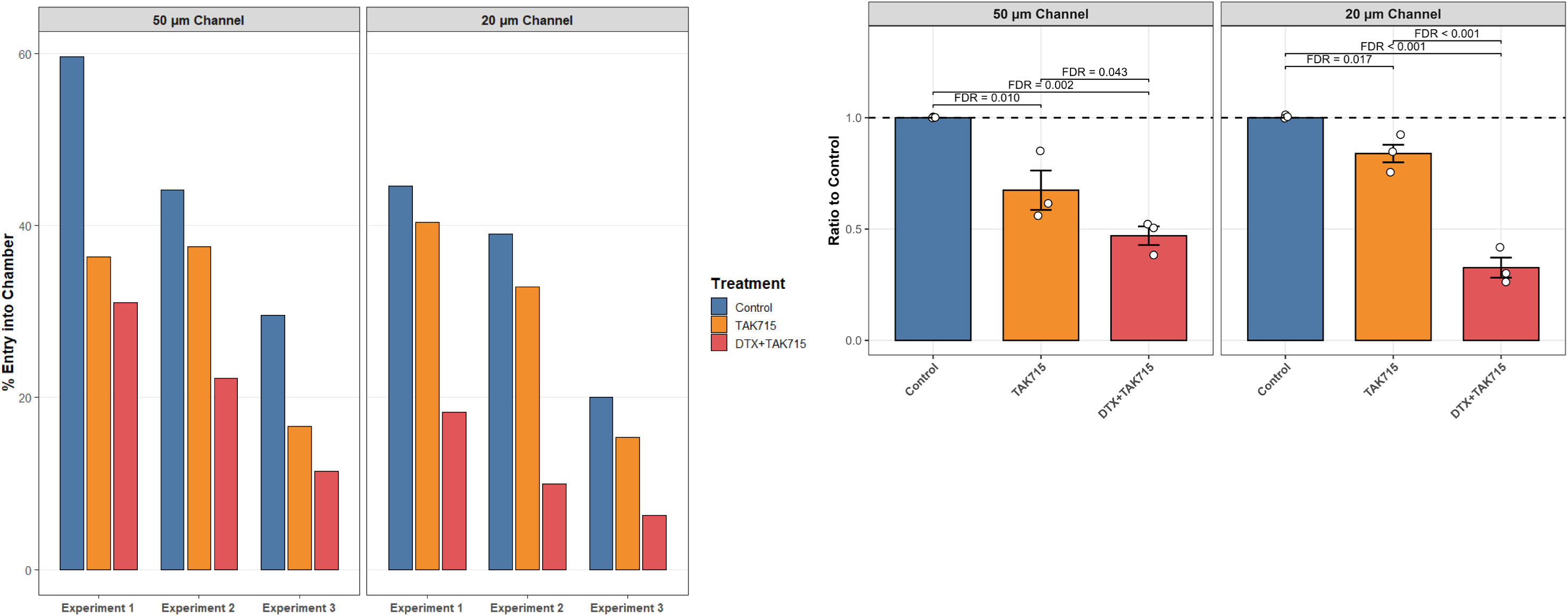
Docetaxel + TAK-715 combination therapy reduces the migratory potential of mCRPC cells. Microfluidics chamber-based Cell migration assay of AR-ve mCRPC cells to evaluate the effect of the drug combination on the metastatic property of the AR-ve mCRPC cells. TAK-715 reduces cell migration and is potentially effective against metastasis and EMT trans differentiation in lethal PCa. Representative plots in DU145 cell lines show that TAK-715 single-agent and combination therapy with DTX reduces the entry of lethal PCa cells into 50 and 20 μm wide μ-channels, indicating a potential role of TAK-715 in abrogating the metastatic potential. i) n=3 experiments; ii) Summary of the results expressed as ratio to the no-treatment control.

**Figure 3B.**
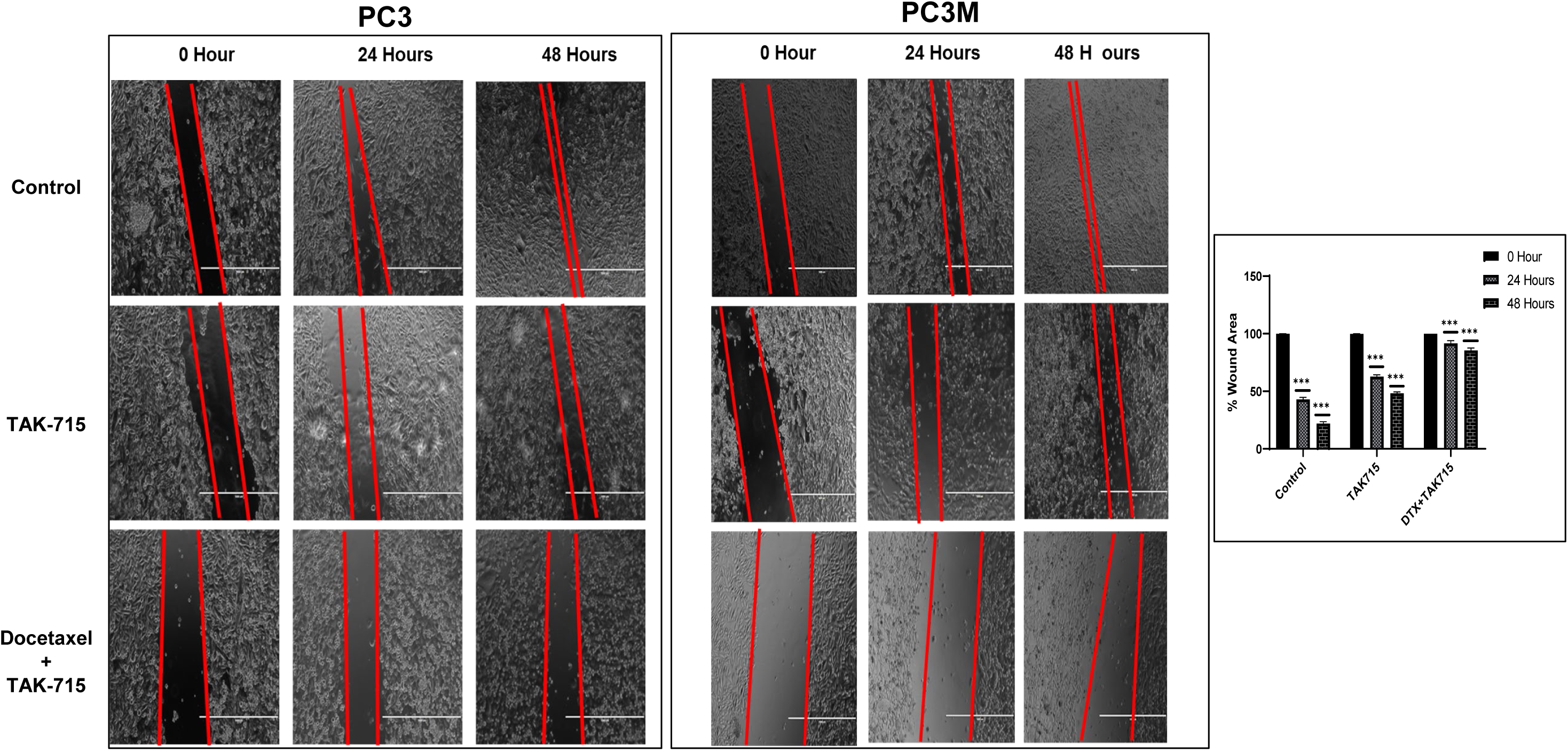
Docetaxel+TAK-715 combination treatment slows down migration of mCRPC cells. Wound Healing Assay of the AR-ve mCRPC cells exposed to TAK-715 single agent and Docetaxel+TAK-715 combination treatment. Representative plots show results of wound healing (Scratch) assay using (A) PC-3 and (B) PC-3M cells. Cell migration after 24 and 48h TAK-715 single-agent and TAK-715+DTX combination was assessed by measuring the scratch size. Images were captured before (0h) and after (24 hours and 48 hours) drug treatments (Significance P-value * = p ≤ 0.05). Bar graphs showed a significant reduction in cell migration (wound healing) following TAK-715-based single-agent and combination treatments.

### Gene expression profiling revealed novel genes and molecular pathways associated with TAK-715 treatment

Next, we performed NGS-based RNA sequencing (bulk RNA-seq) analysis of the untreated, TAK-715-treated, and DTX+TAK-715 combination-treated AR-ve mCRPC cell lines to investigate whole-transcriptome profiles. RNA-seq data were QC-ed, pre-processed, and normalized for differential gene expression analysis. Average quality scores were thoroughly above Q30 for all libraries in both R1 and R2. Heatmaps and Volcano plots were generated to display the differentially expressed genes (DEGs) for each treatment regimen **(Figure 4A-B)**.

**Figure 4A.**
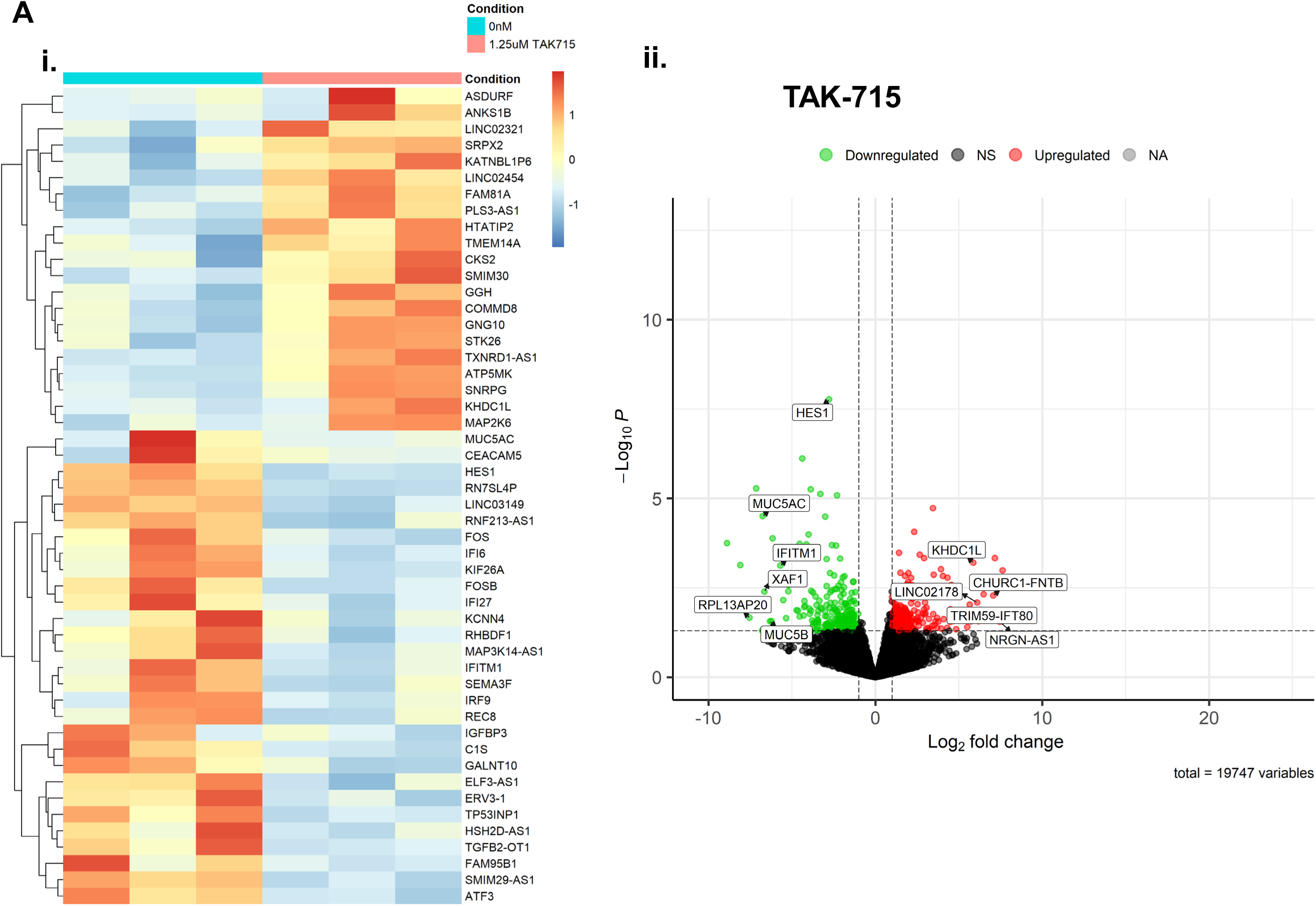
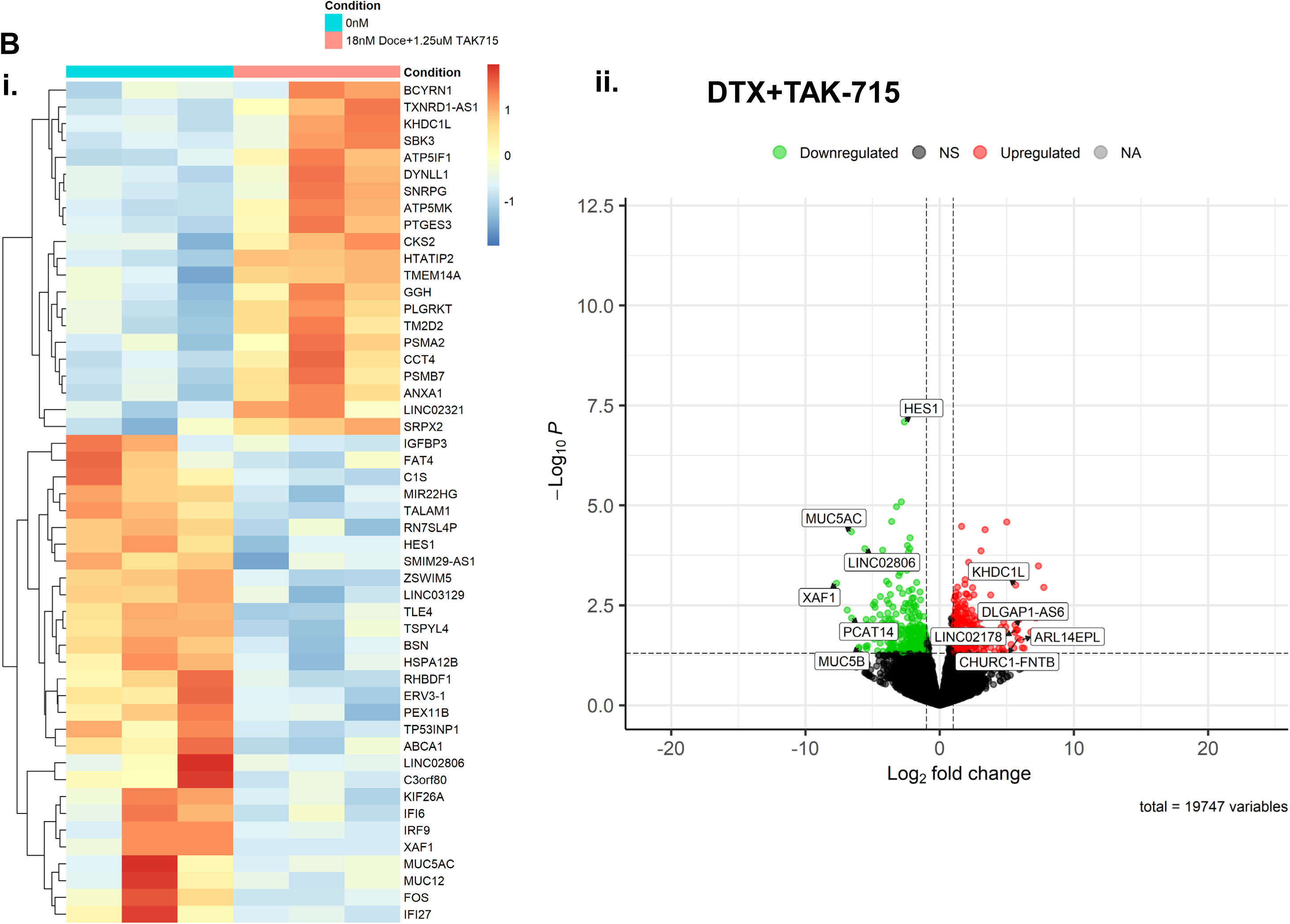
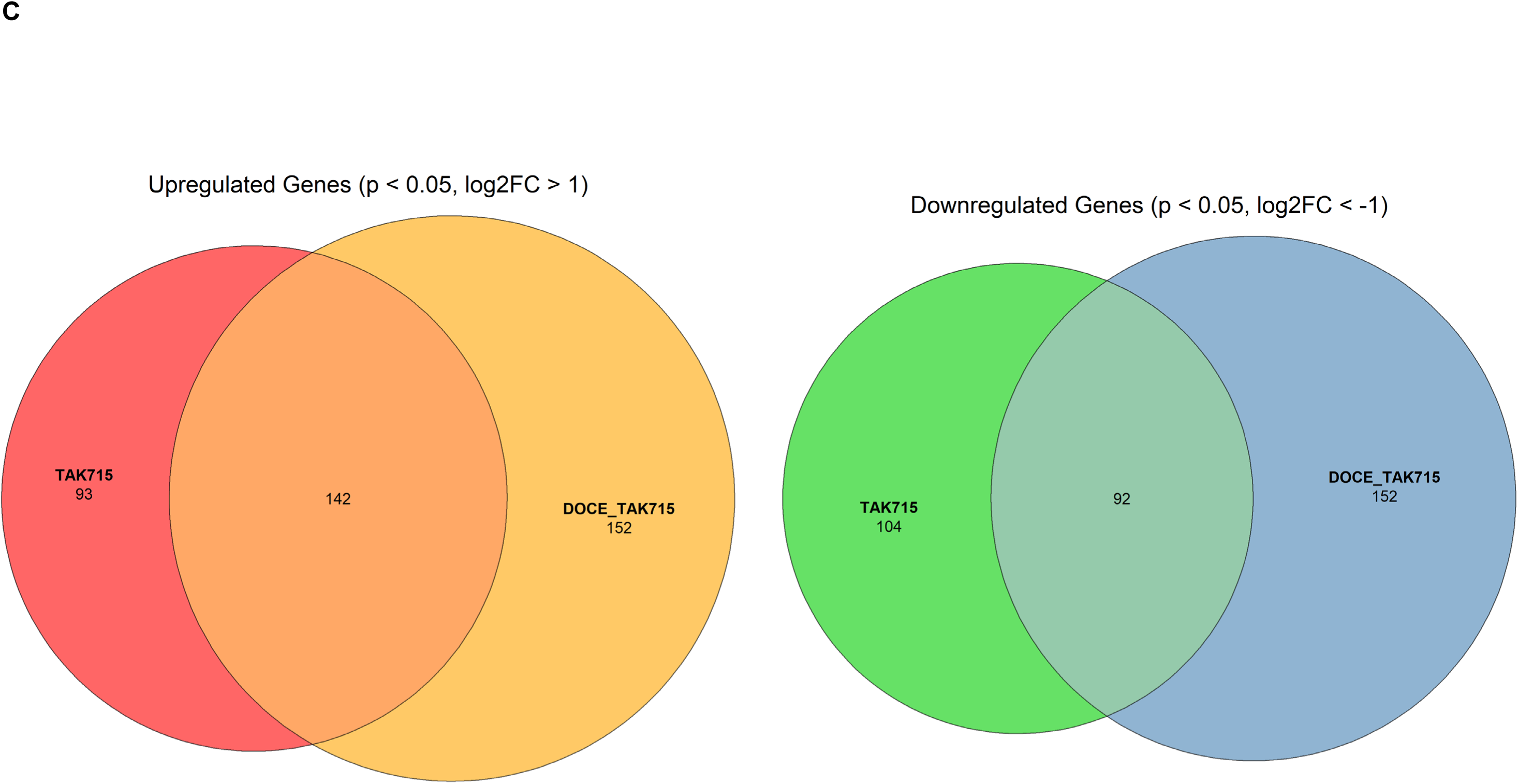
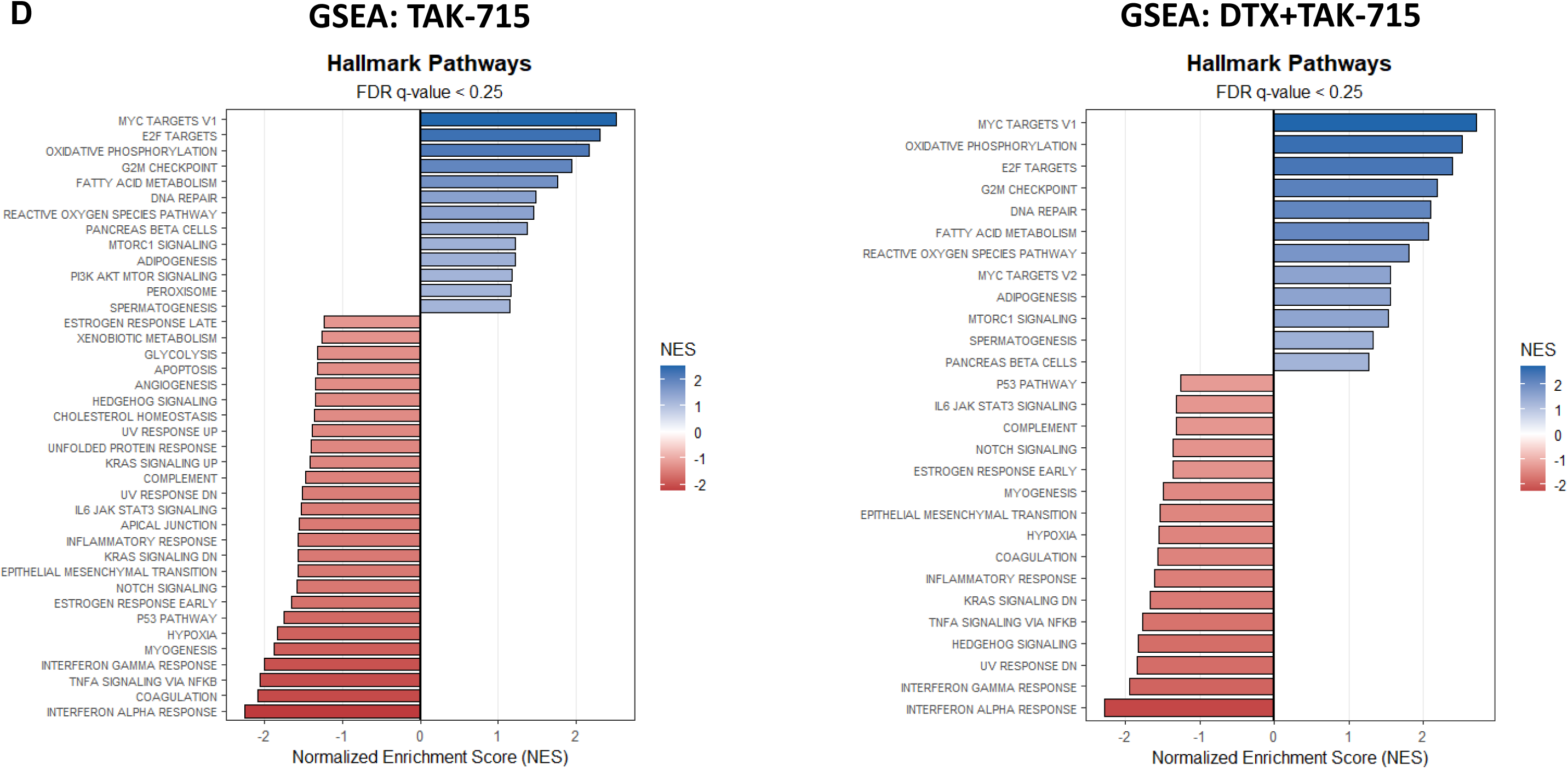
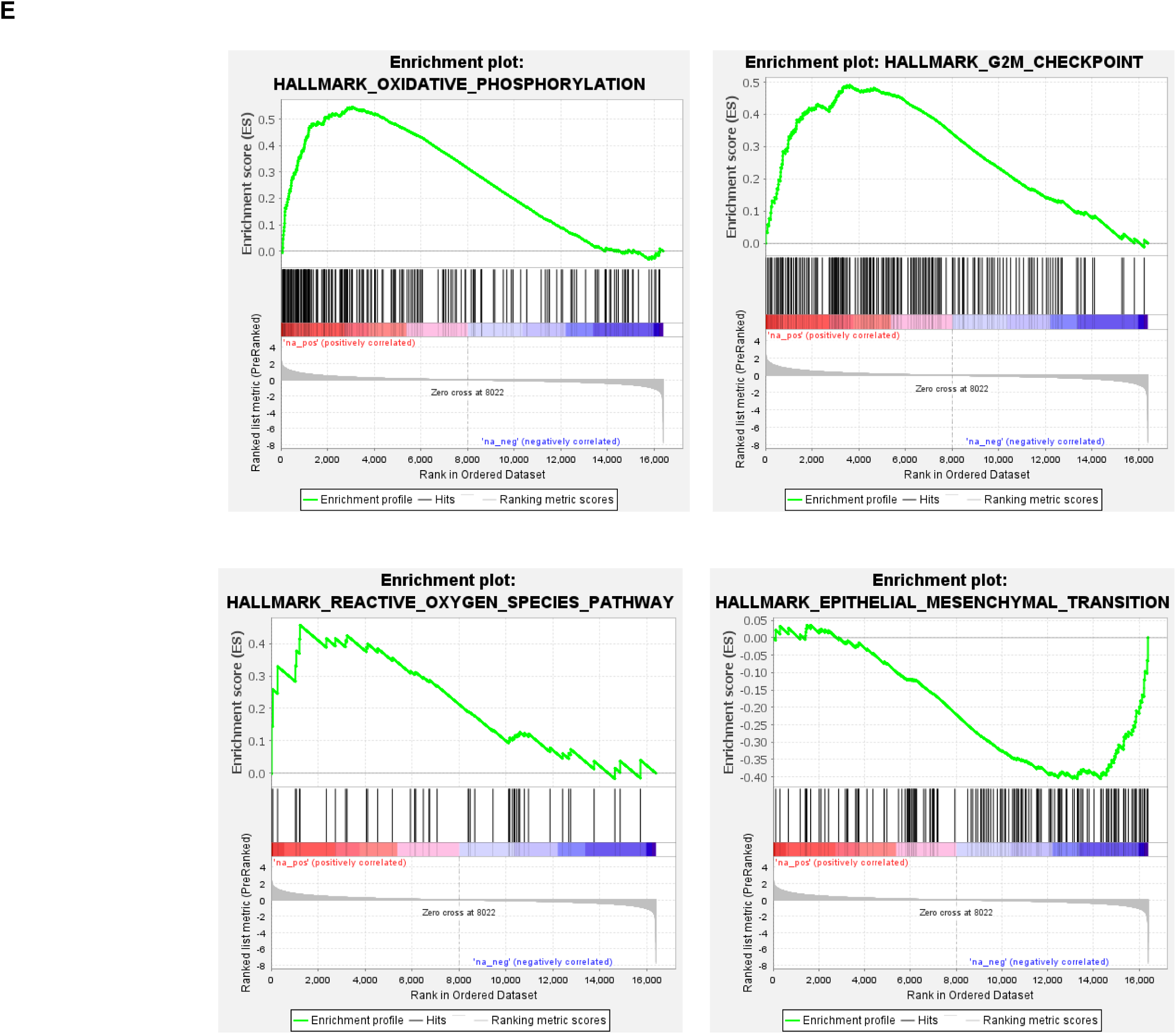
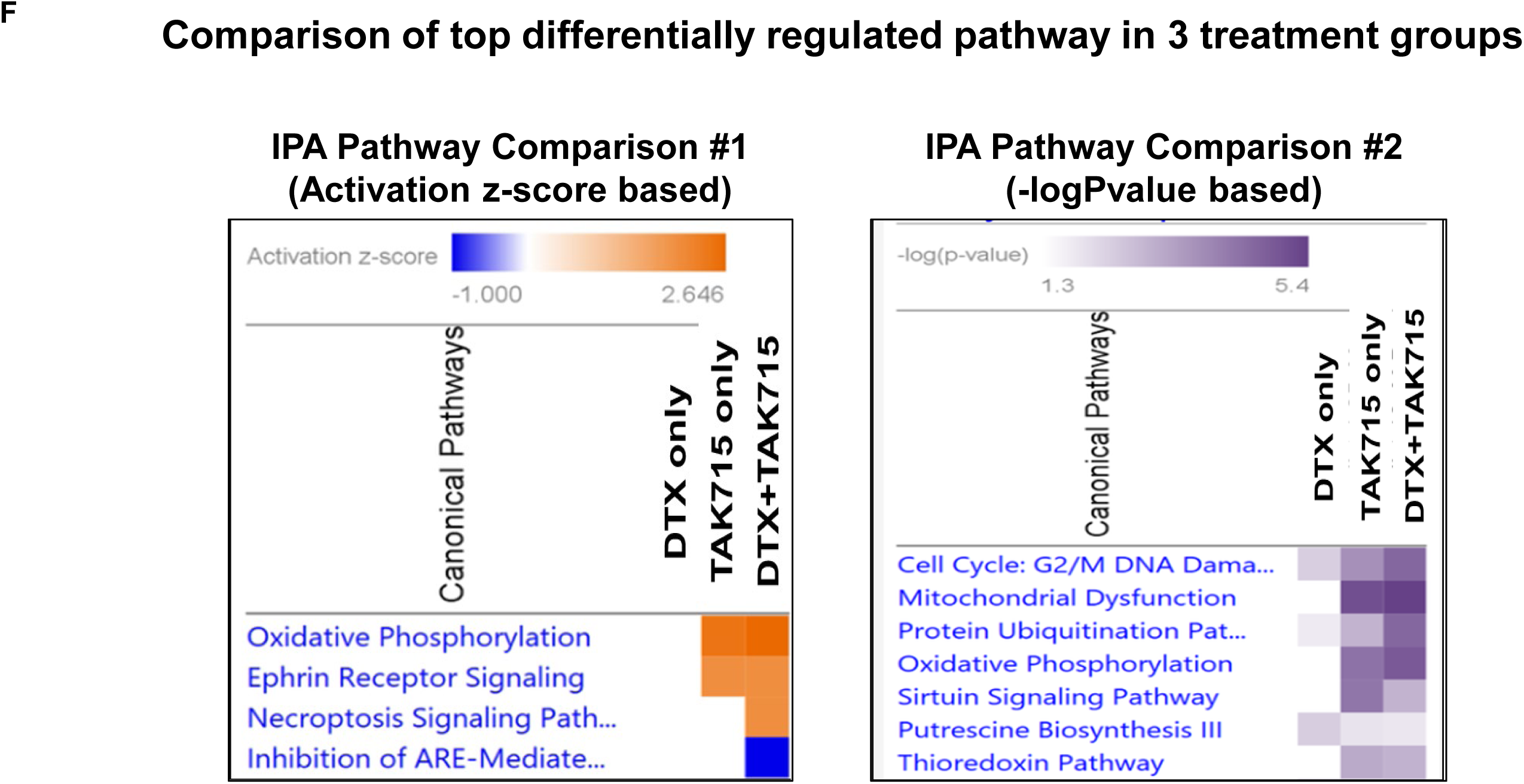
Differential Gene Expression Profile shows top differentially expressed genes in response to the Docetaxel treatment vs Combination Treatment. Differential gene expression profiling analysis results. **A)** TAK-715 single-agent treatment. **B)** DTX+TAK-715 combination treatment. (i) Heatmaps representing top differentially expressed genes (DEGs) following DTX, TAK-715 single-agent, or DTX+TAK-715 combination treatments in AR^null^ mCRPC cell lines (n = 3), 24h following drug exposure. Log2 ratios are depicted in a color scale where red represents upregulation and green represents downregulation. Columns represent cell lines, and rows represent genes. Prior to hierarchical clustering, gene expression values were filtered (samples with max TPM < 1 were removed), and the z score was normalized. (ii) Volcano plots representing differentially expressed genes (DEGs) following DTX, TAK-715 single-agent or combination treatment in human mCRPC cell lines 24 h following drug exposure(|fold-change|> 2, and p < 0.05). Log2 ratios are depicted in a color scale where RED represents upregulation and BLUE represents downregulation. **C)** Euler plots representing unique and common DEGs (p<0.05) between TAK-715 single-agent, and DTX+TAK-715 combination treatments in AR^null^ mCRPC cell lines (n=3). **D-E)** Gene Set Enrichment Analysis (GSEA) showing top pathways enriched in response to treatment. **F)** Ingenuity Pathway Analysis (IPA): A detailed heatmap comparing top differentially regulated pathways for Docetaxel single-agent treatment, TAK-715 single-agent treatment, and DTX+TAK-715 combination-treated cells. IPA analysis confirmed the GSEA results showing mitochondrial dysfunction, oxidative phosphorylation, and cell cycle arrest at the G2/M phase as the top differentially regulated pathways.

Compared to untreated cells, TAK-715 single-agent treatment revealed 431 uniquely differentially expressed genes (p<0.05; fold-difference ≥ 2; Upregulated: 235; Downregulated: 196). Furthermore, 538 genes were differentially expressed following DTX+TAK-715 combination treatment (p<0.05; fold-difference ≥ 2; Upregulated: 294; Downregulated: 244). 234 genes were common between the two lists (Common Upregulated Genes: 142; Common Downregulated Genes: 92) (**Figure 4C**). The top TAK-715-induced down-regulated genes were HES-1, SRSF5, TSPYL2, and TSPYL4, while the top upregulated genes were Cyclin B1, CHAC2, and SSBP1.IPA and GSEA analysis revealed mitochondrial dysfunction, oxidative phosphorylation, cell cycle arrest at G2/M phase, and Epithelial-Mesenchymal Transition (EMT) as the top differentially regulated pathways.

Pathway analysis using Gene set enrichment analysis (GSEA; **Figure 4D-E**) and Ingenuity Pathway Analysis (IPA; **Figure 4F**) based on the DEGs revealed significant enrichment of the following gene sets as the top differentially regulated networks/pathways in response to TAK-715-containing treatments: mitochondrial dysfunction, oxidative phosphorylation, reactive oxygen species, cell cycle arrest at the G_2_/M phase, and Epithelial-Mesenchymal Transition (EMT), in addition to the downregulation of inflammatory and interferon-responsive transcriptional programs like TNFα Signaling via NFκB and IFN-α/β (Type I interferon) (**Figure S4**).

### TAK-715 down-regulates HES1, a transcriptional repressor associated with cancer stemness and multi-drug resistance

Among the top DE genes, HES1 was downregulated in both TAK-715-treated and TAK-715+DTX-treated groups (∼6-7 fold). We further validated the downregulation of the HES1 gene in response to our combination treatment by quantitative real-time PCR. The data show that our drug combinations indeed downregulate HES1 in both taxane-sensitive AR-ve mCRPC cell lines (by ±80%) and taxane-resistant AR-ve mCRPC cell lines (by ±40 %). **(Figure S5A)** We also validated the TAK-715-induced down-regulation of HES-1 at the protein level by western blot analysis. **(Figure S5B)**.

To investigate the clinical implications of HES1 in the context of PCa, we performed *in silico* analysis using the prostate cancer adenocarcinoma database (PRAD) in the Oncomine database and the TCGA portal^18,19^. The Oncomine database suggests HES1 expression is elevated in prostate cancer tissues compared to normal prostate tissues (**Figure 5A**). Analysis of TCGA data showed that HES1 expression is associated with a higher Gleason score, which is a measure of the severity and risk of PCa (**Figure 5B**). Further, expression of HES1 was elevated in metastasis/progression and was significantly higher in nodal metastasis (**Figure 5C**). Next, we validated the dependency of cancer cells on HES-1 for proliferation and survival using the DepMap portal. The DepMap portal is a Broad Institute project to uncover these gene dependencies across hundreds of cancer cell lines ^20^. The distribution of HES1 across cell lines and negative Chronos scores (For example, the Chronos score for HES1 perturbation in the DU145 cell line is -0.178) for HES1 perturbation indicates a strong selective dependence on HES1 for cell survival/proliferation (**Figure 5D**).

**Figure 5.**
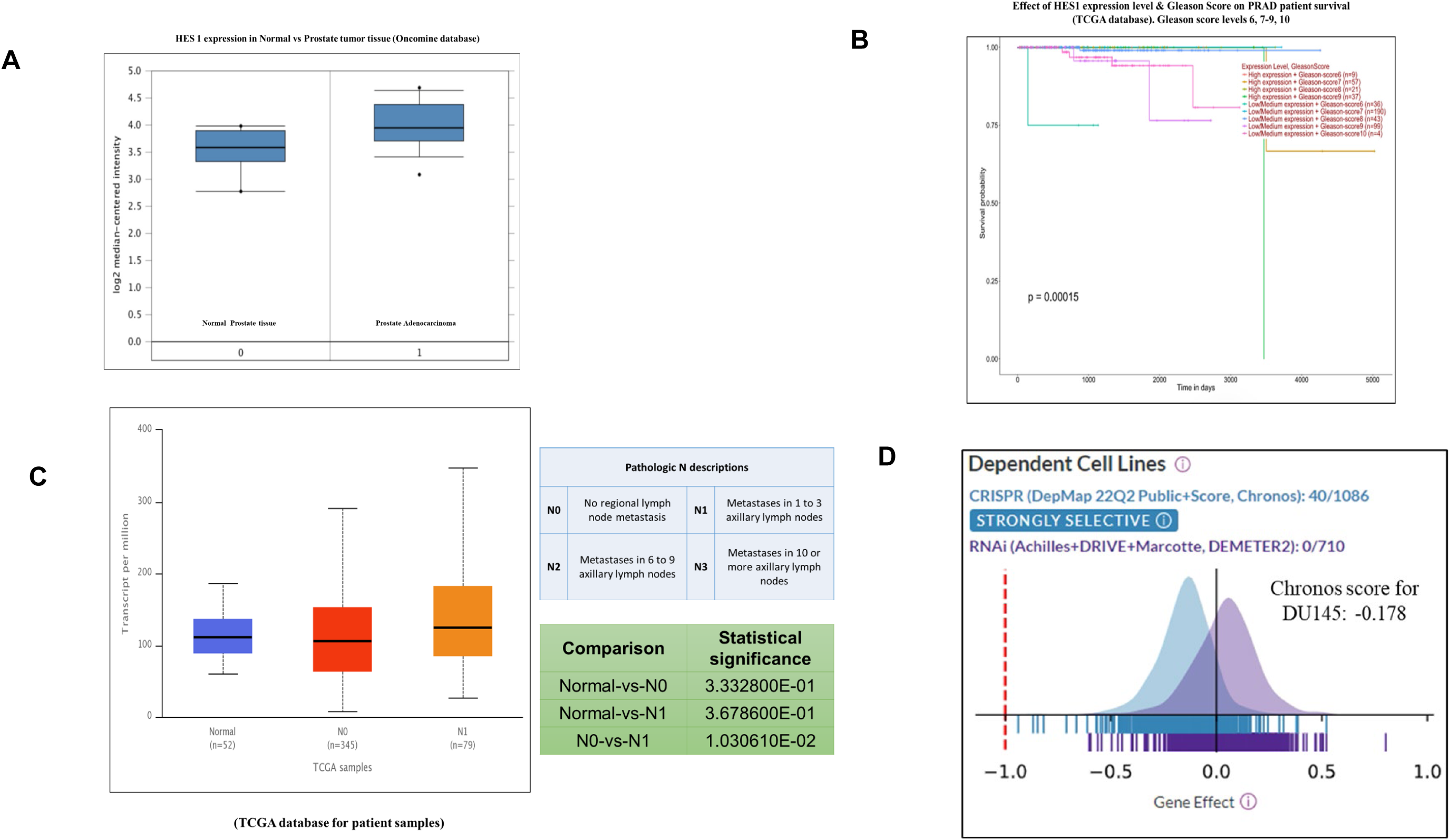
HES1 is a transcriptional repressor associated with cancer stemness and multi-drug resistance. *In silico* analysis of HES1 expression and its impact in prostate cancer. A) HES 1 expression in Normal vs. Prostate tumor tissue (Oncomine database). (B) Effect of HES1 expression level & Gleason Score on PRAD patient survival (TCGA database). Gleason score levels 6, 7-9, 10. (C) Gene expression level of HES1 in PRAD in the different stages of nodal metastasis. (D) DepMap dependency portal data of the the effect of HES-1 expression in PCa cell lines (Chronos score of DU145 is displayed).

### TAK-715 treatment arrests the cells in the G_2_/M phase

We measured the TAK-715 treatment-induced differences in the distribution of cell cycle phases using Propidium Iodide (PI). Cells were treated with TAK-715, either as a single agent or in combination with DTX, for 48 hours, followed by staining with PI and flow cytometry analysis. We observed a significantly higher percentage of the cells that were arrested at G_2_/M phase in response to the combination treatment compared to the untreated and single agent treatment (**Figure 6A**).

**Figure 6A.**
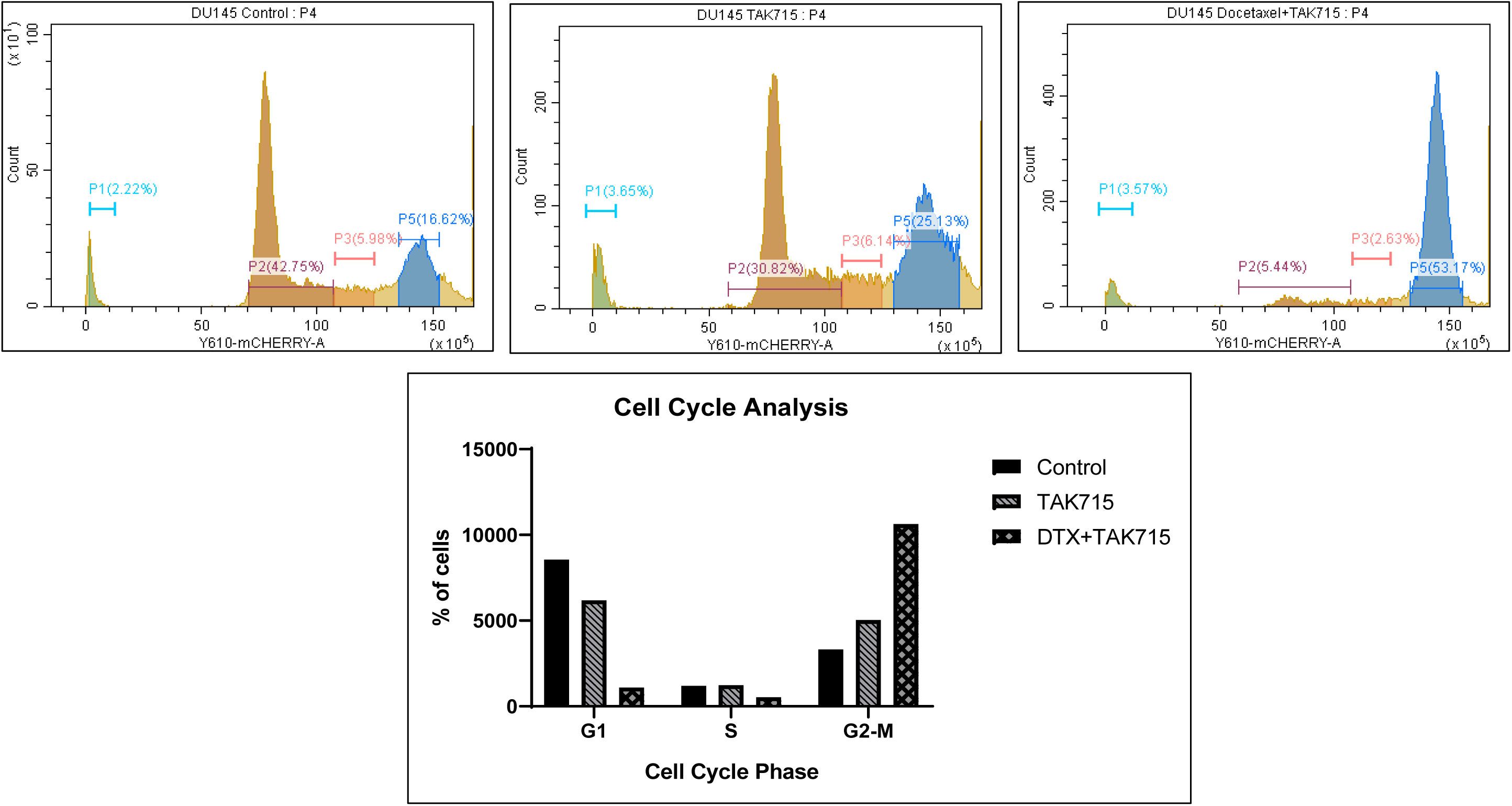
Pathway validation: TX+ TAK-715 combination leads to the cell cycle arrest at G2/M phase in mCRPC cells. Cell Cycle Analysis: Propidium Iodide based Analysis of distribution of different phases of cell cycle in Control (0.5% DMSO), TAK-715 single agent and DTX+TAK-715 combination treated AR-ve mCRPC cells (DU145). Data are presented as the mean ± SEM of three separate experiments (n = 3/study).

### TAK-715 induced elevated DNA damage in AR^-ve^ mCRPC cell lines

Next, to assess the extent of DNA damage, we treated the cells with the DTX+TAK-715 combination for 48 hours and measured the DNA damage induced by TAK-715 treatment using Comet Assay. Undamaged DNA remains in the nucleus, and the damaged, fragmented DNA migrates through the cavity. The shape resembles a comet, with a circular head corresponding to undamaged DNA, and the tail representing the damaged DNA. So, the longer and brighter the tail, the higher the level of damage. The olive moment is (tail length) x (% DNA in tail length). From our comet assay data, we observed that DNA% % in tail, tail length, as well as in Olive moment (DNA% % in tail x tail length) were significantly increased in the DTX+TAK-715 treated cells, which indicates enhanced DNA damage in response to our drug combination (**FIGURE 6B**).

**Figure 6B.**
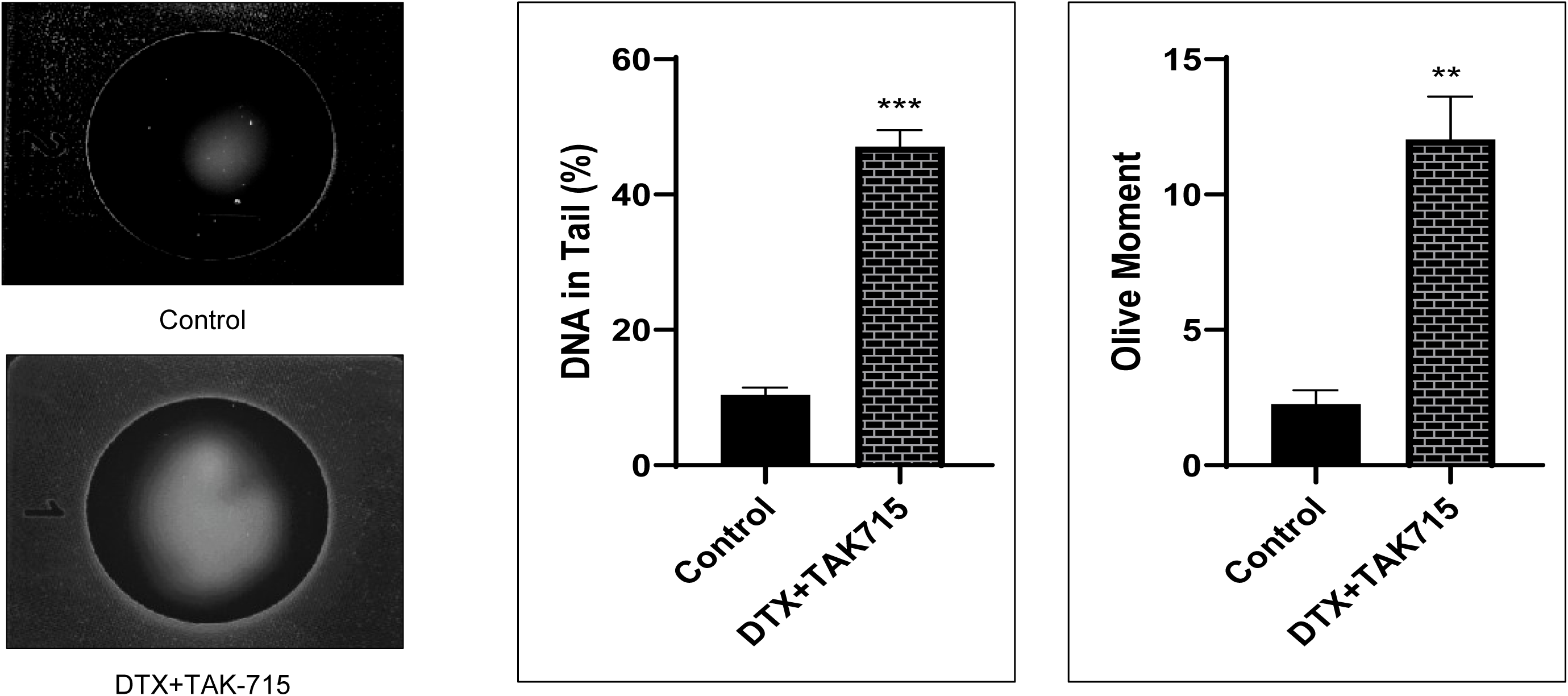
Pathway validation: TX+ TAK-715 combination enhances DNA damage in mCRPC cells. Comet assay to measure the extent of DNA damage in AR-ve mCRPC cells (DU145) post Docetaxel-TAK-715 combination treatment.

### TAK-715 reduced the mitochondrial membrane potential and upregulated cellular ROS generation in AR-ve mCRPC

To investigate if TAK-715 induces its cytotoxic effects through mitochondria-mediated pathway, we measured the mitochondrial membrane potential using JC-1 dye (Abcam). JC-1 is a cationic carbocyanine dye that accumulates in mitochondria. The dye exists as a monomer (green fluorescence) at low concentrations and changes color from green to red in energized mitochondria. The cells were treated with either a single agent (IC_50_ dose of each drug) or in combination (IC_50_ dose of both drugs) for 24 hours. We observed a significant shift from red fluorescence to green fluorescence (**FIGURE 6C**). The decrease in the red/green fluorescence indicated mitochondrial depolarization, which caused the JC-1 dye to become monomers from its aggregate form.

**Figure 6C.**
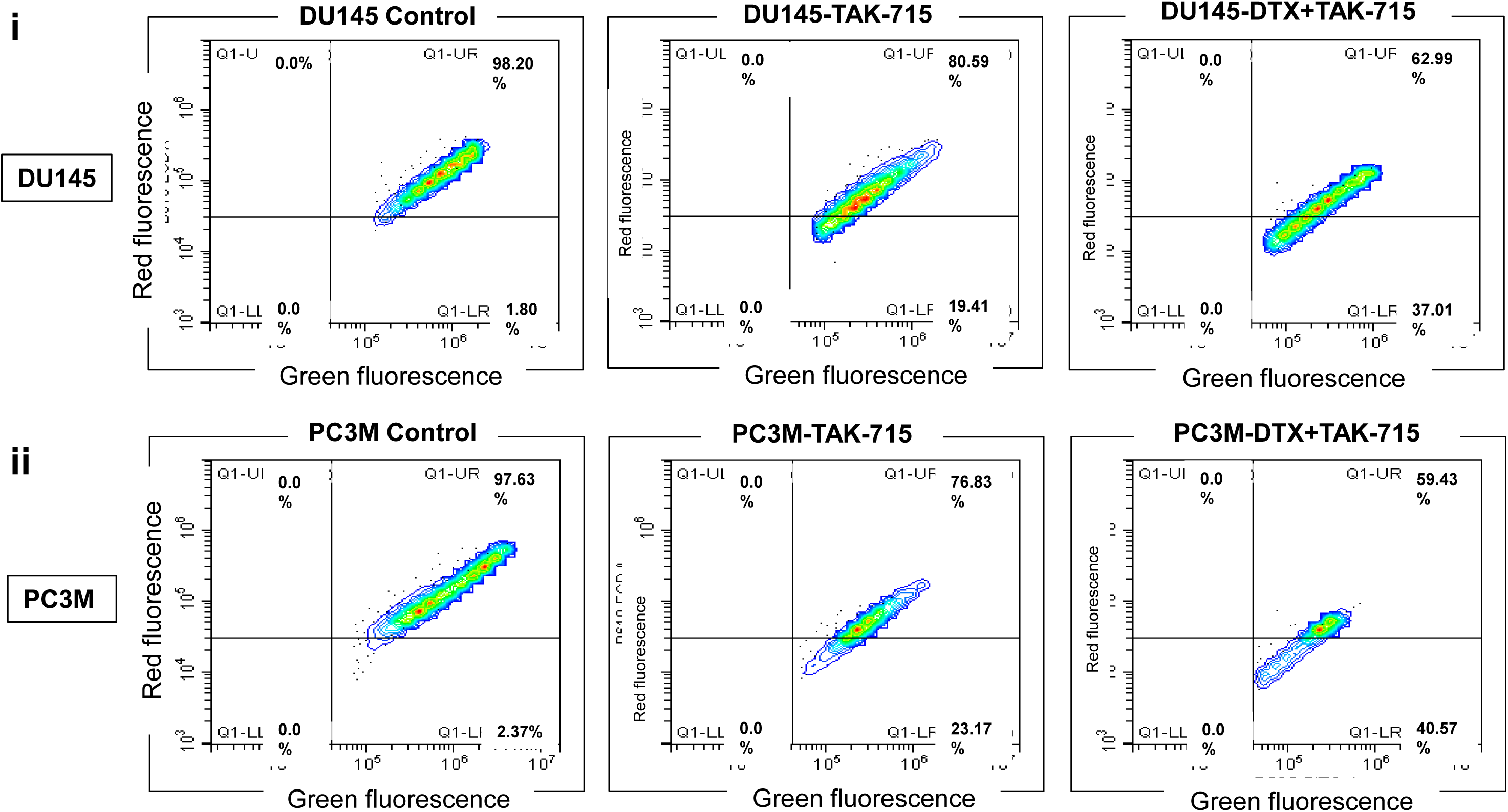
Pathway validation: TX + TAK-715 combination leads to the mitochondrial dysfunction in mCRPC Cell Lines. Mitochondrial membrane potential measurement assay by JC-1 dye accumulation in Control (0.5% DMSO), TAK-715 single agent and DTX+TAK-715 combination treated AR-ve mCRPC cells (i) DU145 (ii) PC3M.

Intracellular reactive oxygen species (ROS) level, a principal factor behind cellular stress-mediated apoptosis, was assessed at different time points using the fluorogenic probe 2,7-dichlorofluorescein diacetate (DCFDA), a cell-permeable non-fluorescent probe that shows fluorescence when it is oxidized. First, we exposed the cells to DCFDA, followed by treatment with TAK-715, either single-agent or combination treatments. Our data indicate time-dependent up-regulation in response to treatments at 2, 4, 8, and 24 hours (**FIGURE 6D**). Moreover, we observed significantly higher levels (p ≤ 0.05) of fluorescence intensity in combination-treated cells compared to single-agent treatment, as indicated by enhanced cellular ROS production. Finally, the DHE assay was performed to confirm TAK-715-induced superoxide generation as an indicator of cellular ROS level in mCRPC cells (**FIGURE 6E**).

**Figure 6D.**
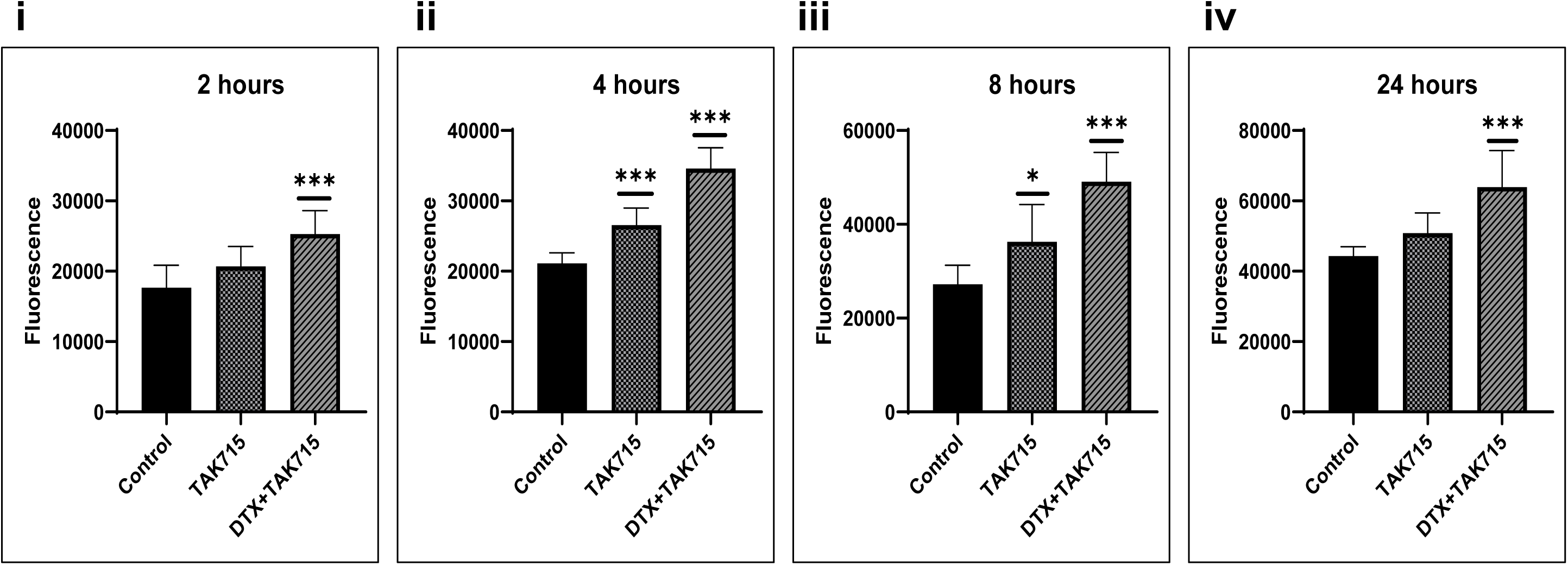
Docetaxel + TAK-715 combination therapy upregulates cellular ROS generation in mCRPC cells. DCFDA assay to measure cellular ROS level in Control (0.5% DMSO), TAK-715 single agent and DTX+TAK-715 combination treated AR-ve mCRPC cells after (i) 2 hours, (ii) 4 hours, (iii) 8 hours, (iv) 24 hours.

**Figure 6E.**
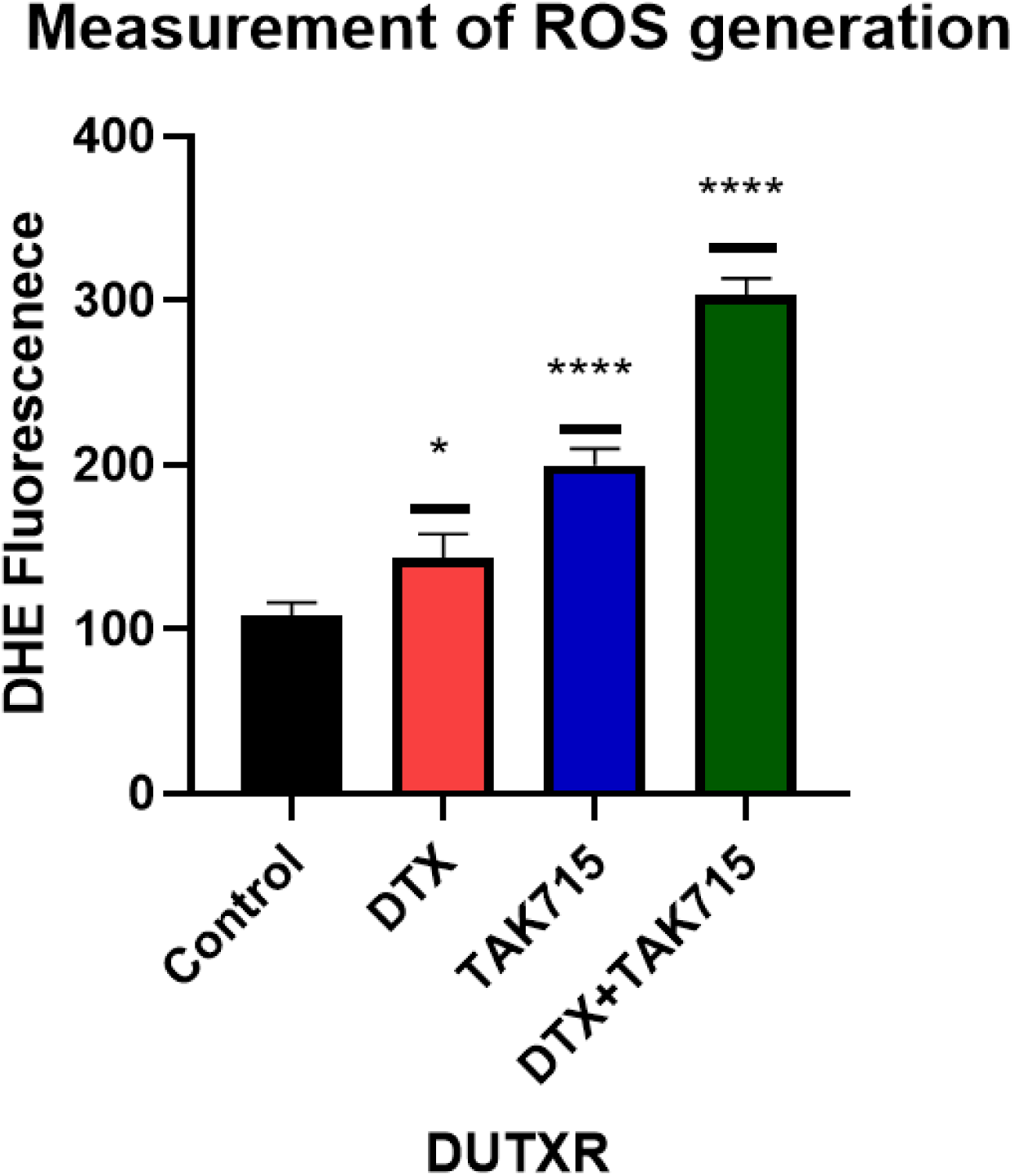
Docetaxel + TAK-715 combination therapy upregulates cellular ROS generation in mCRPC cells. DHE assay to measure cellular ROS level in Control (0.5% DMSO), TAK-715 single agent and DTX+TAK-715 combination treated AR-ve mCRPC cells (DUTXR) after 24 hours.

### TAK-715 reduces the oxygen consumption rate (OCR) in MCL cells

Previous studies have shown that cancer cells have high levels of oxidative phosphorylation, which directly correlates with stemness. To characterize mitochondrial bioenergetics in taxane-resistant AR^-ve^ mCRPC cells (DUTXR) and the effect of TAK-715, we measured the oxygen consumption rate (OCR), which is directly proportional to oxidative phosphorylation, using Seahorse Extracellular Flux Technology. **Figure 6F** shows that TAK-715 was significantly more effective at reducing OCR in DUTXR cells than DTX, suggesting that TAK-715 may indirectly abrogate hypoxia-mediated drug resistance.

**Figure 6F.**
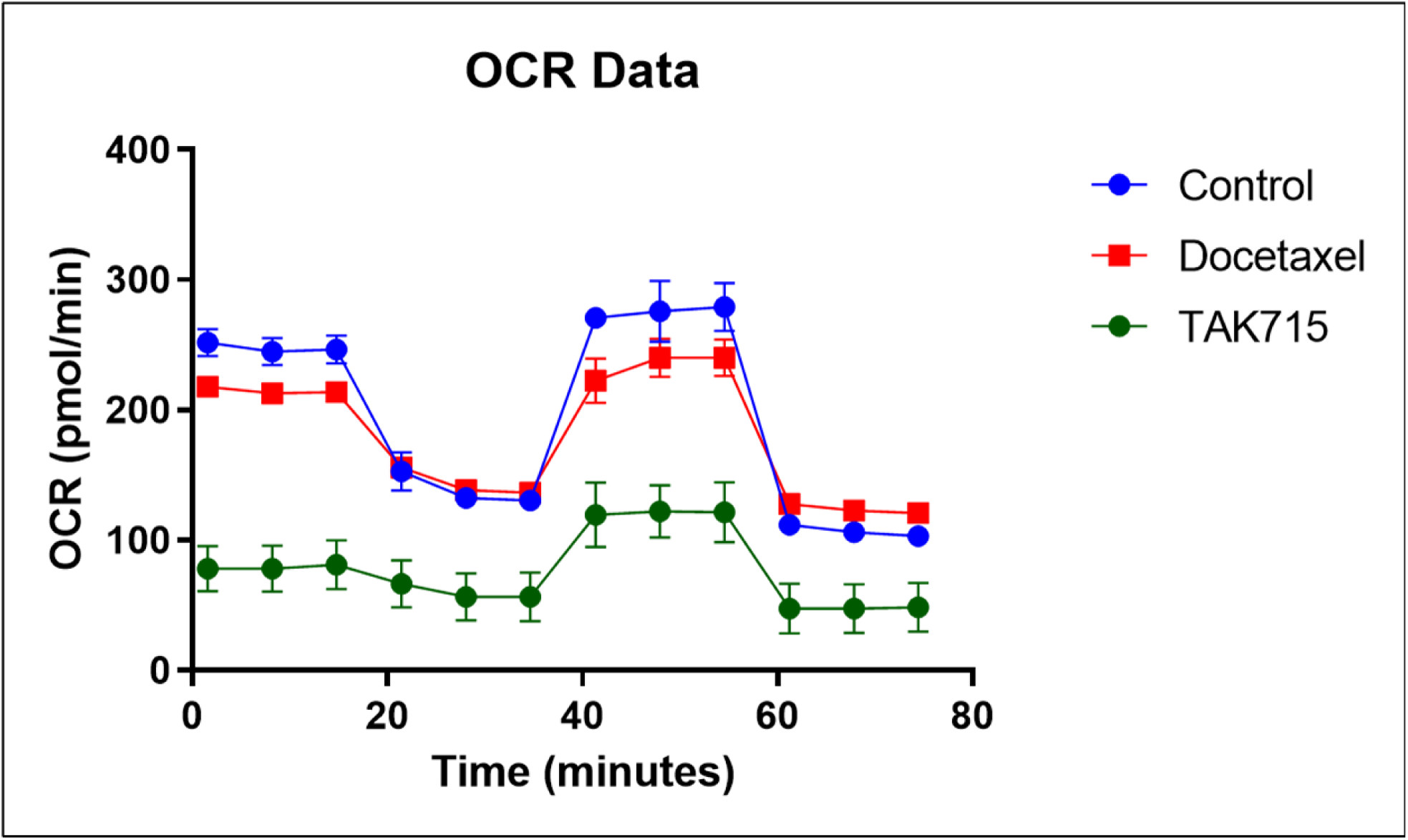
TAK-715 reduces the oxygen consumption rate (OCR) in Prostate Cancer cells. TAK-715 reduces Mitochondrial respiration (measured by Oxygen Consumption rate) in DUTXR cells.

### Single-cell analysis identified TAK-715 treatment-induced changes to subclonal architecture

Pre-vs-Post TAK-715 treatment single-cell transcriptome analysis showed that TAK-715 targets the sub-clones involved in drug resistance and cancer stemness in the taxane-resistant AR-ve mCRPC cell line DUTXR (**Figure 7A**). TAK-715-treated cells show erosion of Clusters 2, 3, 4, and 5 and enhancement of Cluster 6. Pathway analysis of these eroded clusters, based on the expression of the top DE genes, revealed enrichment in molecular networks associated with tumor aggressiveness and disease progression.

**Figure 7A.**
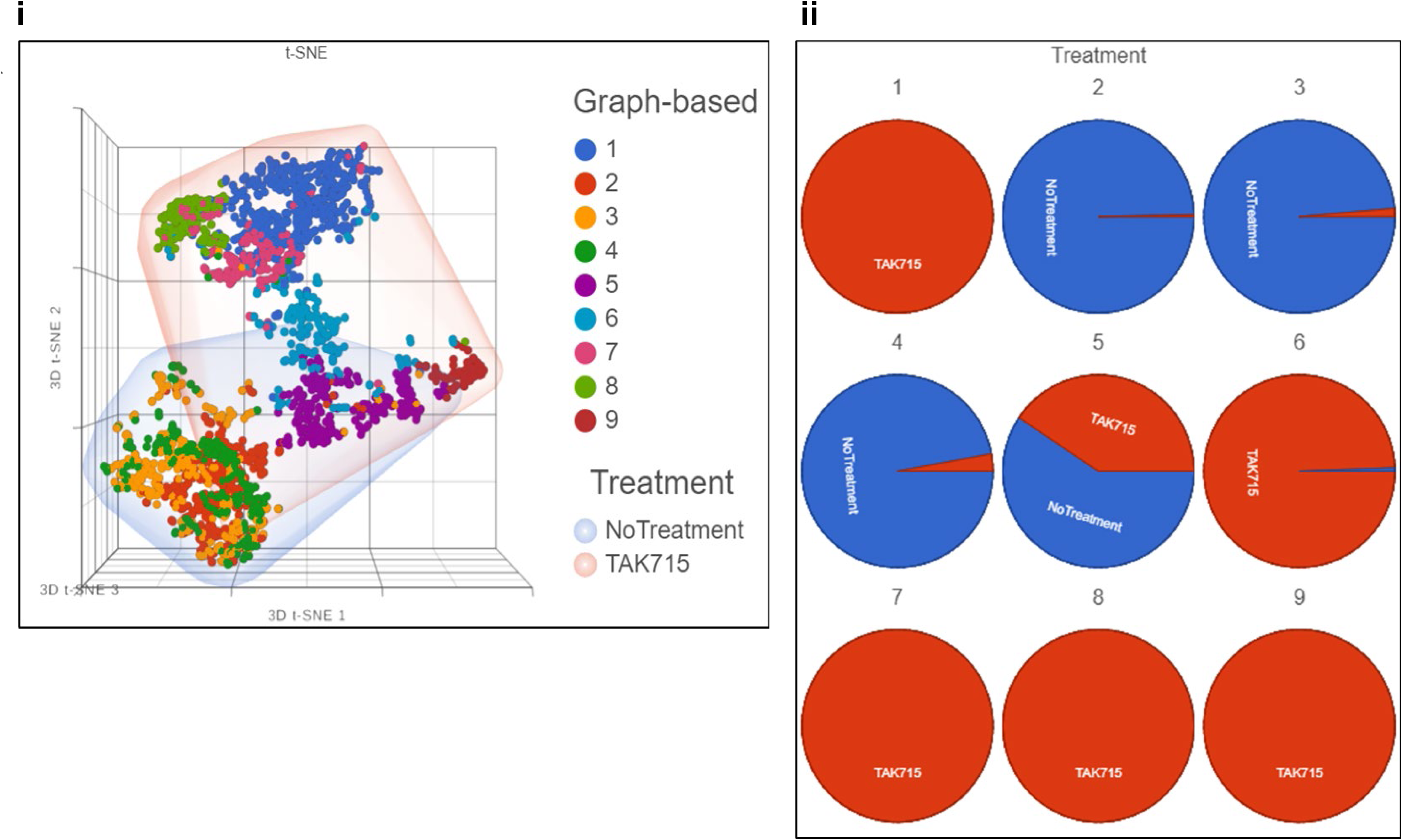
scRNAseq analysis suggests TAK-715 erodes sub-clones responsible for drug resistance and stemness. **i.** tSNE plot showing untreated and TAK7-715-treated DU145 cells. **ii.** TAK-715 treatment leads to the depletion of clusters expressing oncogenes.

### TAK-715 reduces stem cell load in mCRPC lines

#### TAK-715 impaired the Clonogenic properties of mCRPC cells

We investigated the efficacy of our drug combination on the reproductive health of the cells by colony formation assay. Cells were treated with single agent or combination for 24 hours and incubated for 2 weeks, followed by staining with crystal violet. We observed significantly lower numbers of colonies in TAK-715 single-agent and combination-treated wells as compared to no-treatment control (**FIGURE 7B**).

**Figure 7B.**
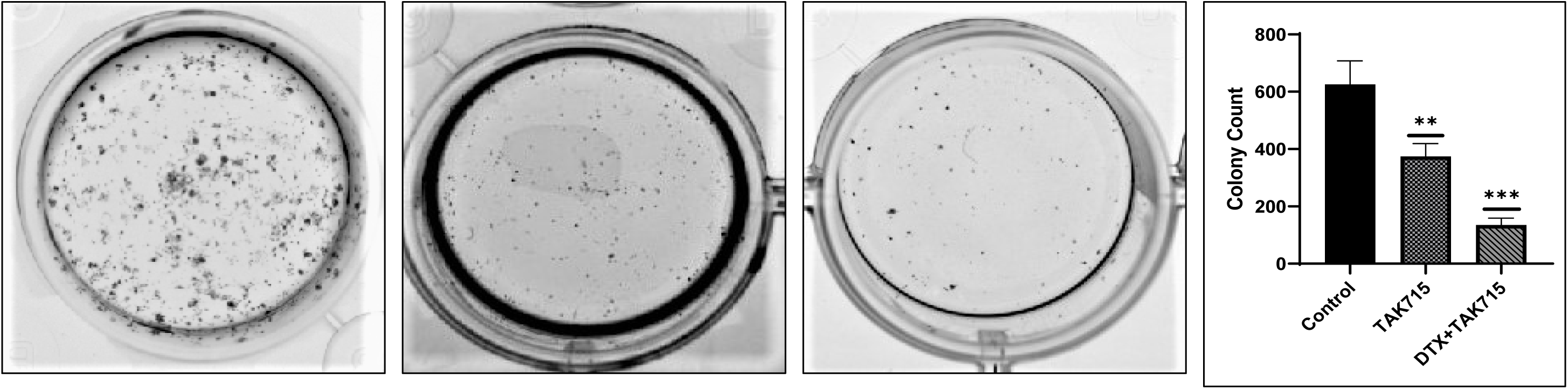
Docetaxel + TAK-715 combination impairs Clonogenic property of mCRPC cells. Effects of Docetaxel+TAK-715 combination on Clonogenic formation of PC3 cells were evaluated by a colony formation assay.

#### TAK-715 diminished the side population load

Side populations are subsets of cells that possess several stem-cell-like features^21^. Briefly, we gated and selected side population (SP) cells from main populations (MP) using DyeCycle violet, pre- and post-TAK-715 treatment. We found that baseline % SP is higher in resistant cells than in parental cells (data not shown). Further, TAK-715 alone or in combination (DTX+TAK-715) reduced SP in taxane-resistant AR^-ve^ mCRPC cells (**Figure 7C**). Our data showed that TAK-715 significantly reduced the side population load (±30%) in taxane-resistant AR^-ve^ mCRPC cell lines.

**Figure 7C.**
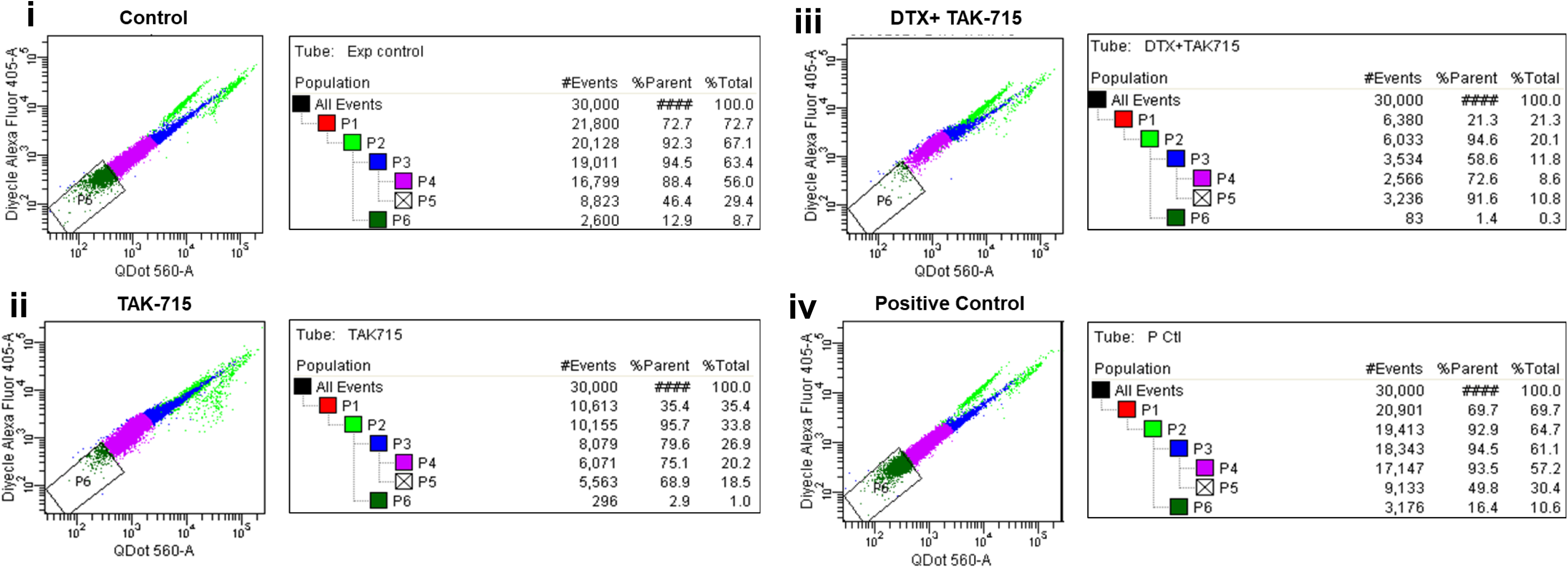
TAK-715 diminished the side population load in taxane-resistant AR^null^ mCRPC cell lines. DyeCycle Violet mediated measurement of side population in taxane-resistant AR^null^ mCRPC cell (DUTXR) post TAK-715 combination treatment. (i) Control (ii) TAK-715 treated (iii) DTX+TAK-715 treated (iv) Positive Control.

#### TAK-715 diminished the markers of ‘cancer stemness’:Cytotoxic effect on the CD44^+^ population

Previous studies reported that in PCa cells, CD44^+^ is a characteristic surface marker for ‘cancer stem cells’ or cancer-initiating cells that play a crucial role in ADT resistance as well as tumor recurrence and metastasis^22^. To study the effect of TAK-715 on this cancer stem-cell population, we first sorted the CD44^+^ population from DUTXR cell lines and treated with two different doses of TAK-715. *In vitro* cytotoxicity assay on these isolated CD44^+^ population cells (**Figure 7D**) shows that TAK-715 was effective in depleting the population of CD44^+^ in a dose-dependent manner. Next, we performed a Caspase 3/7 activity assay on flow-sorted CD44^+^ cells and observed TAK-715 dose-dependent induction of apoptosis in this stem-like cell population compared to the untreated cells.

**Figure 7D:**
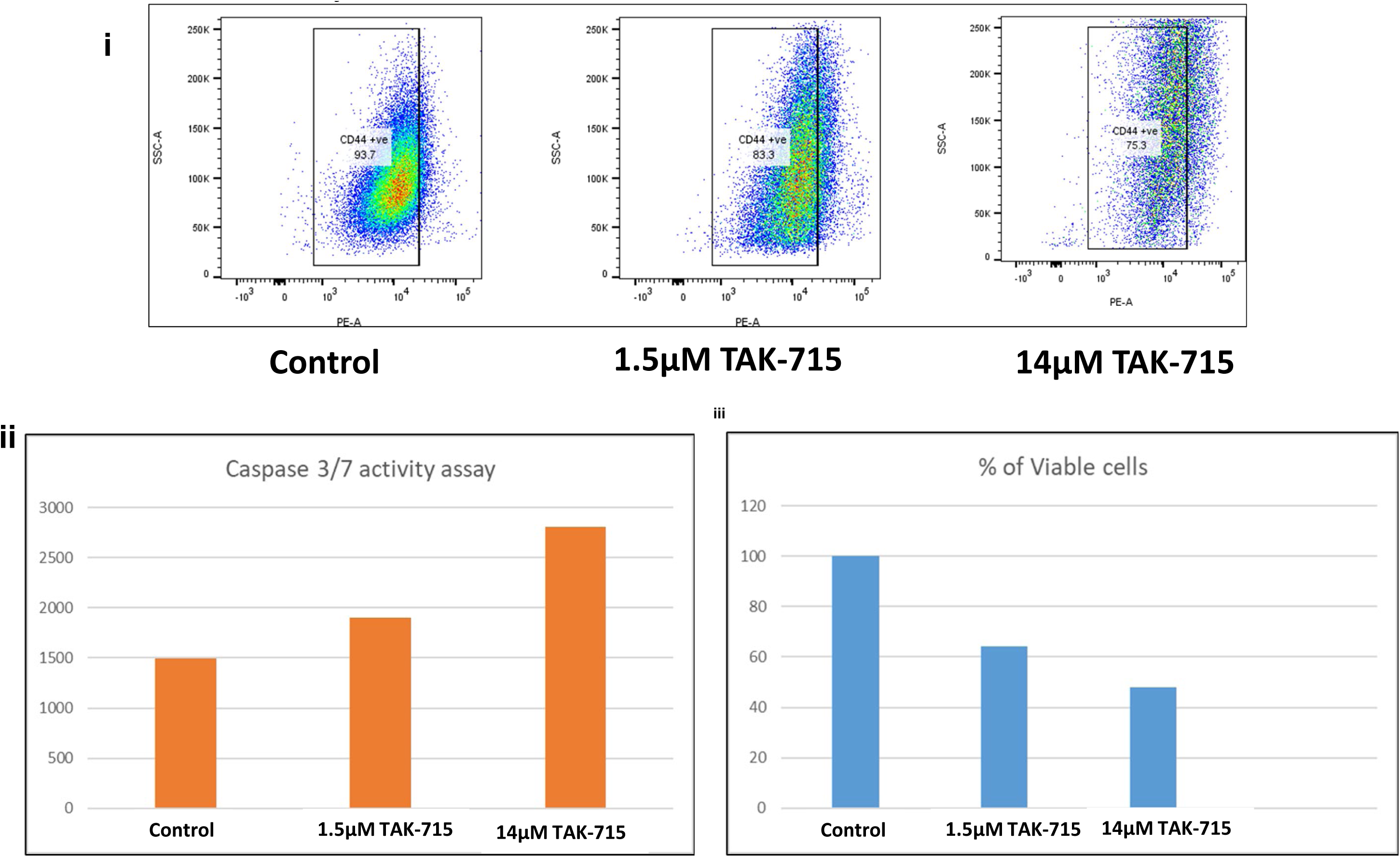
TAK-715 has a significant cytotoxic effect on the CD44^+^ population in AR^null^ mCRPC cells. Effect of TAK-715 on the CD44 population. (i) TAK-715 depletes CD44^+^ population in ARnull mCRPC cells (DUTXR). (ii) TAK-175 induces apoptosis in CD44^+^ cells. (iii) TAK-715 reduced the cell viability of CD44^+^ cells.

### MicroRNA analysis

Causal Network Analysis, a component of IPA advanced analytics, predicted significant upregulation of microRNA-132 and downregulation of miR-21 in response to the TAK-715+DTX combination treatment (**Figure S6A**). The onco-miR, microRNA-132, has several reported anti-tumorigenic activities. *In silico* analysis of multiple PCa datasets showed that the low expression level of miR-132 was associated with poor clinical prognosis, the transition from androgen-dependent (AD) to androgen-independent (AI or AR-ve) stages, and metastasis. miR-21 was significantly upregulated in the GEO PCa datasets (**Figure S6B**). miR-21 is an AR-regulated miRNA that plays a key role in nullifying the effect of castration, driving progression to the AI stage, TX resistance, and cellular invasiveness through down-regulation of tumor suppressor PTEN.

### *In silico* validation of TAK-715 treatment-induced gene signatures using Patient datasets

We applied a novel *in silico* ‘reverse-matching’ approach using TCGA’s prostate adenocarcinoma (PRAD) patient cohort and GSE54460 (106 FFPE radical prostatectomy samples from 100 PCa patients with or without biochemical recurrence (BCR) and DFS data; collected from the Atlanta VA Medical Center, Sunnybrook Research Center (Toronto), and Moffitt Cancer Center) datasets to demonstrate that TAK-715 treatment has the potential to ‘reverse’ differential gene expression signatures associated with PCa lethality ^23^. **PCa patient dataset** (GSE54460; 100 PCa patients with or without biochemical recurrence (BCR) and DFS data): The top genes significantly associated with DFS RNA-seq data from GSE54460 were CKS1B (HR 1.43; p<0.0001; Fold Change_(1vs0)_=-1.34; p=0.003) and DENRP1 (HR 1.29; p<0.0001; Fold Change_(1vs0)_=-1.47; p=0.00013), which were associated with DFS (**Figure 8A**). Our RNAseq data on AR-ve cell lines demonstrated that TAK-715 induced downregulation of CKS1B (FoldChange 2.9; p-value 0.0203), indicating a potential for TAK-715 to treat such advanced PCa patients by ‘reversing’ these signatures.

**Figure 8.**
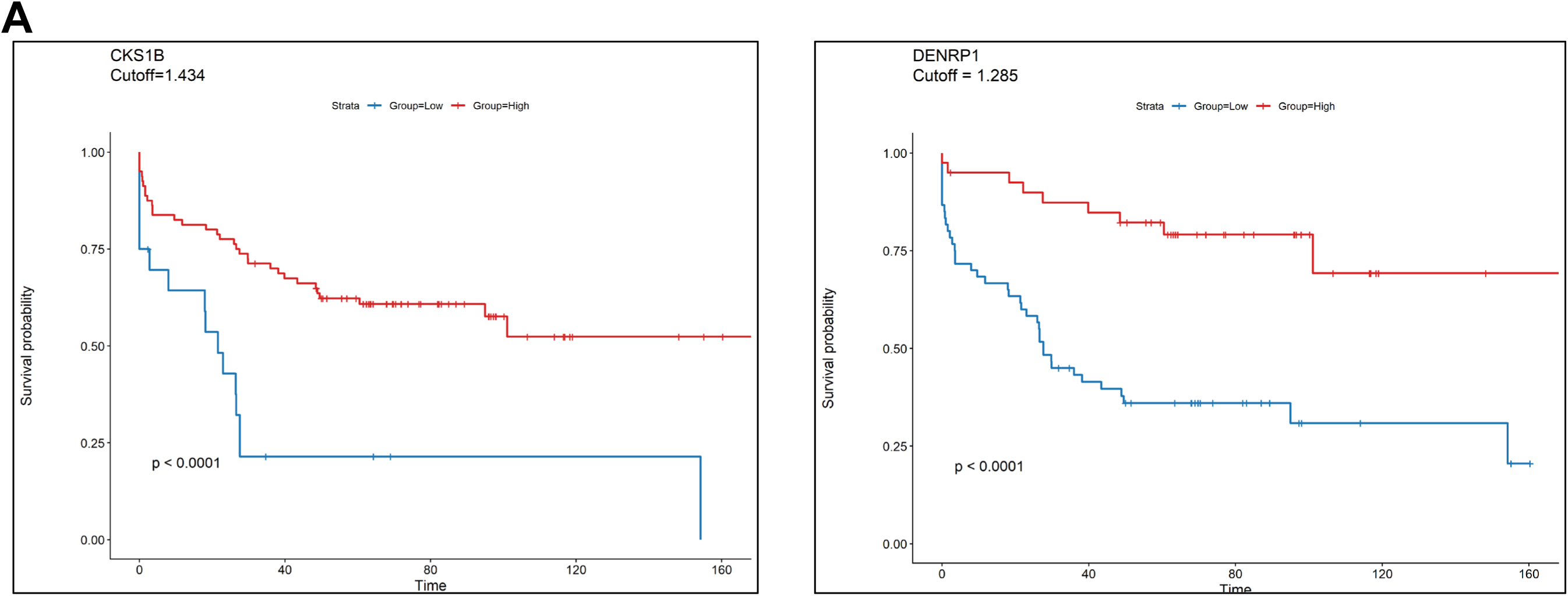
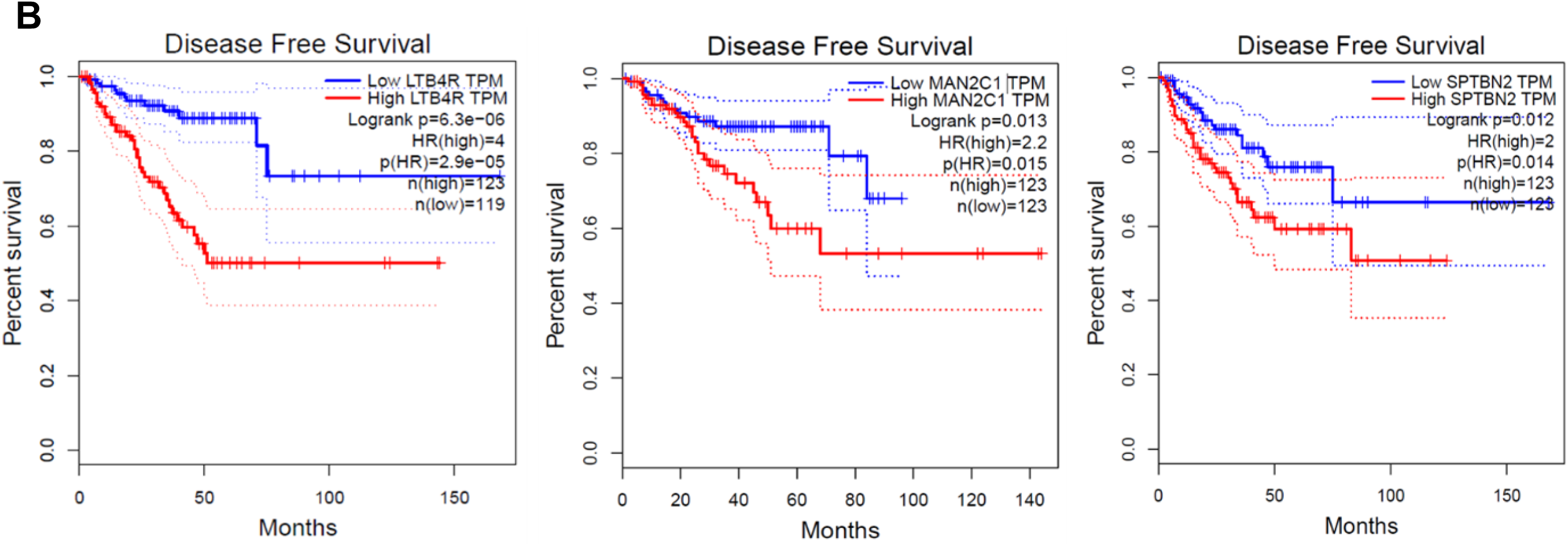
Identification of top TAK-715 treatment-related genes associated with survival in Patient datasets. A. Gene expression data from local PCa tissues from patients enrolled in Atlanta VA Medical Center + Moffitt Cancer Center + Sunnybrook Health Science Center (GSE54460) B. TCGA data on Prostate Adenocarcinoma patients. TAK-715 treatment resulted in significant downregulation of these genes, suggesting potential for TAK-715 to treat such advanced-state PCa patients by ‘reversing’ these gene signatures.

#### TCGA-PRAD dataset

Kaplan-Meier Curves in **Figure 8B** showed that the following genes, which were significantly associated with poor Disease Free Survival in PRAD patients, were downregulated following TAK-715 treatment in PCa cell lines: LTB4R, MAN2C1, and SPTBN2. This demonstrates the potential of TAK-715 to ‘reverse’ these signatures.

## DISCUSSION

Despite improvement in treatment strategies, including first-line and second-line Taxane-based chemotherapeutic drugs (DTX and CBZ, respectively), eventual drug resistance is nearly universal in PCa, with limited therapeutic options^24^. Thus, although the 5-year relative survival in localized and regional PCa is ∼100%, this sharply drops to 37.9% in distant PCa (mCRPC) (NCI SEER Statistics). Therefore, there is an urgent need to formulate new treatment strategies for lethal or advanced mCRPC. We have earlier introduced a pipeline that integrated a pharmacogenomics data-driven approach with a scRNA-seq-based rapid drug screening method and identified the small molecule inhibitor (FK866)-based targeting of NAMPT as a proof-of-concept secondary drug (’secDrugs’) against advanced/lethal PCa, including aggressive, acquired Taxane resistant, and stem-like cell types representing NEPC and EMT phenotypes^14^. Extending the application of the secDrug algorithm, here we show that the inhibition of p38α and CKIδ/€ using TAK-715 is a novel strategy to treat lethal forms of PCa. Further, integrating genome-wide bulk inter-tumor (RNA-seq) and single-cell transcriptomics (scRNA-seq) with cell-based functional assays and patient *in silico* data analysis, we could show the treatment-induced genes and molecular pathways/networks underlying TAK-715 mechanism of action and its potential impact on tumor metastasis, migration, invasion, intracellular ROS activity, and ‘cancer stemness,’ in AR^-ve^ mCRPC cells.

TAK-715 is an orally active small-molecule inhibitor of p38α MAP Kinase that suppresses the release of the proinflammatory cytokines TNF-α and IL-β with almost no inhibitory activity against major CYP enzymes, and was initially developed for the treatment of rheumatoid arthritis^25^. It has been observed that in PCa, both upstream (α-PAK, MEK-6) and downstream (Elk-1, ATF-2) components of the p38 are over-expressed, resulting in enhanced cell proliferation and survival^26^. p38 is involved in IL-6-mediated androgen-independent PCa cell proliferation and phosphorylates and activates key transcription factors (ATF2, Elk-1), which in turn upregulate cell cycle regulators like CCND1 and other genes related to cellular proliferation^26,27^. p38 also inhibits apoptosis through NF-κB activation^28^. Activation of p38MAPK may thus promote aggressive growth of PCa cells and the aberrant AR activity in the absence of androgens, which may promote the onset of androgen independence^29^. Further, hypoxia-associated p38MAPK-mediated AR activation and increased Hif-1α Levels contribute to the emergence of an aggressive phenotype in PCa^29^. Inhibition of p38 decreases IL-1-induced cell proliferation and increases TNF-α-induced cell death^30^. It has also been found that DTX upregulates p53 and p21 in a p38-dependent manner to desensitize PCa cells^31^. Increased p38MAPK activity is associated with DTX resistance in DTX-resistant cell lines^31,32^. Thus, p38MAPK appears to be a potential drug target for mCRPC treatment. Moreover, TAK-715, by cross-reacting with CKIδ/€, also inhibits Wnt/β-catenin signaling, which is involved in cancer cell proliferation and drug resistance in mCRPC^33,34^.

Notably, we observed, in addition to suppression of inflammatory response as previously reported^25^, significant dysregulation of several important genes following TAK-715 treatment, which may be crucial for understanding additional pathways associated with its MOA and MOR. These included downregulation of HES1, RICTOR, and BACH1 and the upregulation of CCNB1, CHAC2, and SSBP1. HES1 is a transcriptional repressor that plays a significant role in cancer stemness, metastasis, antagonism of drug-induced apoptosis, and taxane resistance^35,36^. Knockdown of Hes1 in CD133^+^ positive cells significantly decreases its colony-forming ability, as well as the depletion of the number of cells^35,36^. RICTOR is an essential subunit of the mTORC2 complex that is overexpressed across several cancers and associated with poor survival. RICTOR enhances angiogenesis in PCa, and its downregulation impairs the proliferation of PCa cells^37^. SRSF5 is responsible for the silencing of androgen-inactivating enzymes HSD17B2 and TSPYL2 & TSPYL4, the transcriptional inducer of CYP17A1, which synthesizes DHEA, a major source of intra-tumor androgen^38,39^. Increased expression of cyclin B1 sensitizes PCa cells to apoptosis induced by chemotherapy by decreasing Bcl-2 and increasing p53^40^. CHAC2 acts as a tumor suppressor by inducing mitochondrial apoptosis and autophagy simultaneously through UPR^41^. BACH1 is a transcription factor that is highly expressed in mCRPC cell lines. BACH1 promotes the progression of colorectal cancer, enhances the invasiveness and metastasis in pancreatic and lung cancers, and, by upregulating MMPs, promotes metastasis in PCa^42,43^. On the other hand, NFE2L2/NRF2 is a transcription factor that suppresses PCa cell growth and migration by controlling the transactivation of AR. NRF2 is negatively regulated by BACH1^44^. We observed that TAK-715, in combination with DTX, downregulated BACH and upregulated the expression of NRF-2. In addition, our pathway analysis predicted the stimulation of Ephrin Receptor Signaling and downregulation of OXPHOS and mitochondrial dysfunction following TAK-715 treatment. Ephrin Receptor Signaling is down-regulated in PCa and correlated to higher Gleason scores and shorter survival time^45^. p38α MAPK mediates cell Survival in response to Oxidative Stress via the induction of the redox-sensing transcription factor NRF2 that drives the expression of numerous cytoprotective genes involved in antioxidant response, including superoxide dismutase 1 (SOD-1) and SOD-2^46,47^. Since TAK-715 is a p38α MAPK inhibitor, we observed that it promotes oxidative stress and mitochondrial dysfunction, leading to augmented production of ROS, creating a vicious cycle of mitochondrial damage and oxidative stress that eventually leads to the collapse of the mitochondrial membrane potential, permeabilization of the membrane, and induction of apoptosis.

Furthermore, comparison of our bulk RNA-seq data in PCa cell lines with *in silico* analysis of gene expression data from patients with PCa demonstrated the potential of TAK-715 treatment to ‘reverse’ signatures of poor outcomes/high risk. This includes LTB4R, MAN2C1, and SPTBN2 in TCGA-PRAD and CKS1B in GSE5446 (localized prostate cancer dataset). Interestingly, CKS1B is involved in the disruption of G2/M checkpoint, dysregulation of cell cycle, and abnormal proliferation^48^. Recently, CKS1B was shown to be overexpressed in PCa, promotes cell proliferation and malignant progression mechanisms in PCa^49^. Comparison of pre-vs-post-TAK-715 treatment scRNA-seq data showed the elimination of single-cell Clusters #2 (represented by the genes KRT8, CENPO, NOP16, and Sorcin) and Cluster #4 (PSAT1, RAD21, and CNBP1), as well as the enrichment of genes c-Myc and Cyclin D1. IPO-7, TWF-1, MYOF, FGD6, and MAP4, which have reported oncogenic influences in the cancer cells, including cancer cell growth and proliferation, lethal progression, cell migration, cancer metastasis, EMT, and are also associated with treatment relapse, drug resistance, and poor prognosis^50–55^.

Cancer Metastasis involves dissociation of cancer cells from a primary tumor site, followed by colonization in distant organs by traveling through interstitial tissues, intravasation into the blood or lymphatic vessels, and extravasation into a new site^56^. Thus, cell motility through confining pores plays a pivotal role in the process of metastatic dissemination, during which cells undergo EMT^57^. Therefore, in this study, we used a highly sophisticated microfluidic device-based assay called Confined Cell migration assays as a physiologically relevant model of metastasis, in which we recapitulated diverse microenvironmental cues encountered by cancer cells during locomotion to demonstrate the potential effects of TAK-715 on cancer cell invasion, cell motility, and metastasis^17^.

Since TAK-715 has been shown to be well-tolerated in animal models^25^, we propose conducting *in vivo* validation studies using mouse xenograft models to establish TAK-715 as a promising drug re-purposing candidate for anti-PCa therapy.

## MATERIALS AND METHODS

### *In silico* secondary drug prediction (secDrug)

The proof of the secDrug algorithm is non-trivial and mathematically involved, as reported previously^14^. As data source, we used the GDSC1000 (Genomics of Drug Sensitivity in Cancer; a large-scale pharmacogenomics database of dose-response results (IC_50_ or AUC) on 565 drugs covering a wide range of targets and processes involved in cancer biology in >1000 cell lines representing a wide spectrum of human cancers^58^. For the purpose of this study, we used cell lines (n=136) representing genitourinary cancers (including PCa) that are aggressive and resistant to DTX therapy.

### Drugs & Reagents

Details on reagents, drugs, and antibodies used are provided in **Supplementary Table S1**. All the drugs were dissolved in dimethyl sulfoxide (DMSO) and stored at -20°C.

### Prostate Cancer Cell Lines

AR-ve mCRPC/NEPC cell lines (PC3, PC3M, DU145) were obtained from the American Type Culture Collection (ATCC) (Manassas, VA, USA). The taxane-resistant cell lines PC3-TXR and DUTXR were generated using dose-escalation of taxanes over time (**Figure S7**), as described earlier^14^. The cell lines were authenticated at the source and tested randomly at regular intervals for tissue specimen provenance and cell lineage at the AU Center for Pharmacogenomics and Single-Cell Omics (AUPharmGx) using the GenePrint 24 System (Promega). All cell lines are mycoplasma negative. PC-3 and PC-3M cells were maintained in 10% (v/v) FBS-supplemented F-12K, and DU145 in Eagle’s Minimum Essential Medium (EMEM). PC3-TXR and DUTXR were maintained in RPMI-1640 media with 1% Penicillin-Streptomycin at 37°C, 21% O2, and 5% CO2 in a humidified cell culture chamber (Heracell™ VIOS 160i CO_2_; Thermo-Fisher Scientific™).

### *In vitro* cytotoxicity assays and drug synergy analysis

*In vitro* chemo-sensitivity assays were performed on human PCa cell lines using the MTT (3-(4,5-dimethylthiazol-2-yl)-2,5-diphenyltetrazolium bromide reagent) assay. Briefly, cells were plated in a 96-well culture plate at 2×10^3^ cells/well, incubated for 24 h at 37°C with 5% CO_2_, and then treated with increasing concentrations of DTX, CBZ, or TAK-715 as a single agent, or in combination (DTX+TAK-715 or CBZ+TAK-715). Following 48-hour incubation, the tetrazolium dye MTT was added according to the manufacturer’s instructions, and absorbance was assessed at 550 nm using the Synergy Neo2 Microplate Reader (BioTek, USA) as a measure of mitochondrial enzyme activity. Percent change relative to untreated controls was calculated at each drug concentration, and the effect of drug exposure was determined by constructing cytotoxicity (growth) curves. Half-maximal inhibitory drug concentration (IC_50_) values were estimated by nonlinear regression using a sigmoidal dose-response equation (variable slope). Drug synergy was calculated by comparing single-agent and combination drug-response data based on Chou-Talalay’s combination index (CI) method and the isobologram algorithm (CompuSyn software; Biosoft, US)^59^. CI values between 0.9-0.3 and 0.3-0.1 signify synergism and strong synergism, respectively, between the drugs treated in combination.

### *In Vitro* Live Cell Imaging

Live-cell imaging was performed in phase-contrast mode to directly visualize the cytotoxic effect on cellular morphology and total cell counts.

#### Cellular morphology

PCa cells were seeded at 0.025 × 10^6 cells/ml in 6-well plates and exposed to TAK-715, either as a single agent or in combination with DTX, for 48 h. Three areas with approximately equal cell densities were identified in each well, and images were captured with an EVOS FL digital cell imaging system (Thermo Fisher Scientific, Inc.) using a 10X objective.

For **<u>nuclear morphology</u>**, PCa cells were plated on glass coverslips (1.5×10 ^5 cells/ml), incubated overnight, and treated with vehicle, TAK-715 alone, or TAK-715 in combination with DTX. After 48h, the cells were labeled with NucBlue Live reagent and incubated for 20 minutes. Images were captured using a Nikon Eclipse Ti2 microscope and recorded in bright-field and phase-contrast modes at 20X and 40X magnifications. Images representative of each treatment condition were analyzed using ImageJ software (National Institutes of Health, Bethesda, MD, USA).

### Assessment of cellular apoptosis

#### Annexin V and propidium iodide (PI) staining

Cellular apoptosis was determined by the Annexin V-PI staining. Briefly, cells were seeded in 6-well plates at the indicated concentrations and exposed to DTX and TAK-715 as single agents and in combination. After 48h, cells were labeled with binding buffer containing annexin V-FITC (25 µg/ml) and PI (25 µg/ml) as well as 10 mM HEPES, 140 mM NaCl, 5 mM KCl, 1 mM MgCl_2,_ and 1.8 mM CaCl_2_ (pH = 7.4), incubated for 10 min, followed by three washes in binding buffer. Both detached and attached cells were combined, and staining was quantified using a Becton Dickinson FACS Calibur flow cytometer (BD Biosciences, San Jose, CA) at 10,000 events per measurement. The data was analyzed using CytExpert software.

#### Caspase 3/7 activity Assay

Cell death by apoptosis was measured using the Caspase-Glo 3/7 luminescent assay kit according to the manufacturer’s instructions (Promega, Madison, WI). Briefly, 2×10^3 cells/well were seeded into 96-well plates (triplicate) and treated at the estimated single-agent and combination IC50 values, as determined by MTT assay. Following 48 hours of incubation, Caspase-Glo 3/7 reagent was added and incubated for 2 hours, and luminescence was measured using a Synergy 2 Hybrid Multi-Mode Microplate Reader (BioTek, USA). The caspase activity at each concentration was normalized to the untreated controls, and the area under the relative caspase activity curve was calculated using the trapezoidal method in GraphPad Prism (LaJolla, CA, USA).

### Cell Cycle Analysis

The effect of the drug treatments on the cell cycle distribution was assessed by quantifying DNA content using PI/RNase staining solution. Control (no drug) and post-treated cells were prepared for cell cycle analysis by staining with PI (50 µg/ml) in sample buffer [PBS + 1% (w/v) glucose], containing RNase A (100 units/ml) for 30 min at room temperature and analyzed by flow cytometry using a Becton Dickinson FACS Calibur flow cytometer (BD Biosciences, San Jose, CA). Cell cycle data were analyzed using CytExpert (Beckman Coulter Inc., Indianapolis, IN).

### Z’ LYTE Assay

The Z’ LYTE Assay was performed at Thermo Fisher Scientific to identify the targets of TAK-715. Briefly, 100 nL of 100X Test Compound (for each test concentration) in 100% DMSO was taken in a black 384-well plate. 2.4 µL of Kinase buffer, 5 µL of 2X Peptide/Kinase Mixture (for each of the target kinases), and 2.5 µL of 4X ATP Solution were added to each well, followed by a 30-second plate shake. The plate was incubated at room temperature for 60 minutes for the Kinase Reaction. 5 µL of Development Reagent Solution was added, followed by a 30-second plate shake. The plate was incubated again for 60 minutes at room temperature for the Development Reaction. The fluorescence reading was captured in a plate reader, and the data were analyzed.

### Side Population

The effect of TAK-715 on the side population cells was investigated using DyeCycle Violet (cell-permeable DNA binding dye) (Thermo Fisher Scientific, Waltham, MA, USA). Briefly, 1×10^6/ml cells were cultured in 6-well plates and treated with TAK-715 at the indicated concentrations. After 24 h, cells were stained with 5μM Vybrant® DyeCycle Violet and 1μg of 7-AAD (to gate out dead cells) for 30min at 37°C and immediately analyzed using a Becton Dickinson FACS Calibur BD LSR II flow cytometer (BD Biosciences, San Jose, CA) at 10,000 events per measurement.

### Isolation of the CD44^+^ population

DUTXR cells were collected and washed with PBS, followed by permeabilization using cold methanol. Cells were then washed twice with PBS and resuspended in 500 μL of antibody dilution buffer containing CD44PE-conjugated antibody, followed by a 1-hour incubation at room temperature in the dark. The cells were washed with PBS and sorted using the MoFlo XPD Flow Cytometer. The sorted cells were immediately put in culture using DMEM/F12 (1:1) basal media containing human epidermal growth factor, basic fibroblast growth factor, and recombinant human leukemia inhibitory factor.

### Colony formation assay

Colony formation assay was performed to measure the effect of the drug treatments on the proliferative capacity of mCRPC cell lines. Briefly, cells were seeded in a 6-well plate at 0.025*10^6^ cells/ml, incubated overnight, and treated with vehicle (0.5% DMSO), DTX single agent, TAK-715 single agent, and a combination of DTX and TAK-715. The cells were then harvested and plated in a 24-well plate at a concentration of 1000 cells/well and incubated for 1-2 weeks at 37 °C. The colonies were fixed with 100% methanol and stained with 0.5% Crystal Violet staining solution. Images were captured using an EVOS FL digital cell imaging system (Thermo Fisher Scientific, Inc). Images were recorded in bright-field and phase-contrast modes at 20X and 40X magnifications and analyzed using ImageJ software (National Institutes of Health, Bethesda, MD, USA).

### Cell migration/Scratch Assay

Wound Healing Assay is a standard and commonly used method to study cell migration/motility^60^. Briefly, Cells were plated in 6-well plates at 1×10 ^5 cells/well and incubated for 48 h to reach 95% confluency. The monolayer was scratched with a SPLScar Scratcher 6-well Tip at a width of 0.50 mm at the center of the well. TAK-715, as a single agent or DTX+TAK-715 combination, was applied to the cells in the respective wells. F-12K culture medium supplemented with 10% FBS containing the vehicle (0.05 % DMSO) was added to the cells in the control wells. Micrographs of the wound areas were obtained at 0, 24, and 48 hours using an EVOS FL digital cell imaging system (Thermo-Fisher Scientific, Inc.). Images were recorded in brightfield and phase contrast modes at 20X and 40X magnifications. The area of the initial wound (at 0 h) and the “gap area” were measured at 48 hours using ImageJ software (National Institutes of Health, Bethesda, MD, USA).

### Microfluidic (μ)-channel Cell Migration Assay

The fabrication of the Polydimethylsiloxane (PDMS)-based μ-channel assay using standard multilayer photolithography and replica molding has been demonstrated earlier^17^. The device consists of an array of parallel channels of variable width (3-50 μm), fixed length (200 μm), and height (10 μm). Perpendicular to the μ-channels are two larger, 2D-like channels that served as cell seeding and chemoattractant inlet lines. Prior to cell seeding, the μ-fluidic devices were coated with 20 μg/mL rat tail collagen type I (Corning) for 1 hour at 37 ^°^C to facilitate cell adhesion. This allowed us to study the effect of our drug on tumor cell motility through μ-channels of dimensions that mimic the size of channel-like tracks encountered by migrating cells *in vivo*. In this study, PCa cells were seeded in 6-well plates and exposed to TAK-715 as a single agent and TAK-715+DTX combinations at indicated concentrations. Next, 1-1.5 ×10^^5^ cells were introduced into the cell seeding inlet line of the microfluidic channel via pressure-driven flow and were allowed to adhere for 30 min at 37 °C, 5% CO2. The cell suspension was removed and substituted with a serum-free medium. Medium supplemented with 10% FBS was added to the chemoattractant inlet line to trigger cell entry into the channels. The devices were placed on an automated Nikon Ti2 Inverted Microscope equipped with a Tokai Stage-Top incubator unit, which maintained cells at 37 °C and 5% CO_2_. Cell entry into the channels was recorded via time-lapse microscopy. Images were recorded every 20 min for 10 hours using a 10x/0.45 NA Ph1 objective. The migration efficiency of drug-treated mCRPC cells compared to control was assessed by calculating the percentage of cell entry into the channels (defined by the total number of cells entering the channels divided by the total number of cells seeded within 50μm diameter from the μ-channel entrances as well as cell velocity (defined as net cell displacement divided by the total time of cell tracking). Because our PCa cells did not frequently enter narrower microchannels (≤10 μm), we focused our analysis on wider channels (≥20 μm).

### Mitochondrial membrane potential (MMP) measurement

PCa cells were seeded and treated with DTX, TAK-715, and DTX+TAK-715 for 24 h. Cells were harvested, washed with PBS, and incubated with 100 μL/well of 2μM JC-1 dye for 10 min in the dark at 37°C. Fluorescence was acquired using a Synergy Neo2 Hybrid Multi-Mode Microplate Reader (BioTek, Winooski, VT, USA) at 535 nm. Data were analyzed using CytExpert software by Beckman Coulter (Brea, CA, USA) to determine the change in red/green fluorescence signal ratio, corresponding to the JC-1 monomer/JC-1 dimer ratio, which indicates healthy versus depolarized mitochondria.

### Total cellular reactive oxygen species (ROS) measurement

To investigate the effect of TAK-715 on oxidative stress, we quantified the intracellular ROS levels between pre- and post-treatment conditions. Briefly, cells were plated in black flat-bottom 96-well plates at a seeding density of 2000 cells/well and incubated overnight at 37°C in 5% CO_2_. The next day, cells were washed and incubated with 100μl of 10 μM fluorogenic probe 2,7-dichlorofluorescein diacetate (DCFDA) solution in the dark for 45 minutes at 37°C. The DCFDA solution was then discarded, and cells were washed twice with phosphate-buffered saline (PBS), treated with either vehicle (0.5% DMSO), TAK-715 single agent, or TAK-715+DTX combination at the indicated drug concentrations for 2, 4, 8, and 24 hours. ROS production was evaluated by measuring fluorescence intensity on a Synergy Neo2 Hybrid Multi-Mode Microplate Reader (BioTek, Winooski, VT, USA) at excitation of 485nm and emission of 535nm in endpoint mode.

To measure superoxide levels, PCa cells were pre-incubated with 5 μM DHE for 15 min in the dark at 37 °C and treated with TAK-715-based regimens for 24h. Cells were then washed once with a cell-based assay buffer, and red fluorescence was recorded by the Synergy Neo2 Microplate Reader (BioTek, USA).

### Measurement of Oxygen Consumption Rate (OCR)

We measured the OCR using Agilent Seahorse Extracellular Flux (XF) Technology. Briefly, DUTXR cells were plated in an XFp plate and treated with vehicle control (0.5% DMSO), DTX, and TAK-715 for 24 hrs. Following 24hr incubation, the mitochondrial function was measured by the XFp Seahorse analyzer using the Agilent Seahorse XF Cell Mito Stress Test kit. First, oligomycin and Fluoro-carbonyl cyanide phenylhydrazone (FCCP) were injected sequentially, followed by a third injection of a mixture of Rotenone and Antimycin A. Oligomycin inhibits ATP synthase and reduces OCR, followed by FCCP, which raises OCR to the maximal rate by collapsing the inner membrane gradient and increasing the electron transport chain activity. Lastly, rotenone and antimycin A, which are complex I and antimycin complex III inhibitors, respectively, inhibit the electron transport chain and reduce the OCR to a minimal value. Data were normalized to the protein concentration at the end of each experiment, and graphs were plotted using Agilent Seahorse Wave Desktop software and report generator, MS Excel, and GraphPad Prism.

### Comet Assay

The extent of drug-induced DNA damage was assessed using Trevigen’s alkaline comet assay kit following the manufacturer’s protocol (R&D Systems). Briefly, PCa cells were plated in a 6-well plate and treated with DTX+TAK-715 combination for 48 hours. Cells were then collected, washed with PBS, mixed with low-melting agarose, and immobilized with molten agarose (LMagaore)+ Ethidium Bromide at a 1:10 (v/v) ratio and immediately spread on the CometSlide. The slides were kept at 4°C in the dark for 10 minutes, immersed in ice-cold lysis solution at 4°C for 45 minutes to break open the cell membrane, and DNA was denatured in freshly prepared alkaline unwinding solution (pH>13) for 20 minutes at room temperature. Next, the slides were placed in a cold electrophoresis tank, and alkaline electrophoresis was performed for 30 minutes. Cells were then stained with propidium iodide, and Images were captured using the Gel Doc EZ Gel Documentation System with the OpenComet plugin in ImageJ software (National Institutes of Health, Bethesda, MD, USA).

### RNA Isolation and RNA Sequencing

The effects of DTX and TAK-715, as single agents and in combination, on gene expression in PCa cell lines were investigated using next-generation RNA sequencing of bulk tumor cells. Cells were incubated overnight in a 6-well plate, followed by treatment with TAK-715 single-agent and DTX+TAK-715. After 24 hours, pre- and post-exposure tumor cells were harvested, and high-quality RNA was extracted using QIAshredder and RNeasy kit (Qiagen) and stored at -80°C. RNA concentration and integrity were assessed using a NanoDrop 8000 (Thermo Fisher Scientific, USA) and an Agilent 2100 Bioanalyzer (Applied Biosystems, Carlsbad, CA, USA). An RNA integrity number (RIN) threshold >8 was applied, and RNA-seq libraries were constructed using Illumina TruSeq RNA Sample Preparation kit v2. Libraries were then size-selected to generate ∼200 bp inserts, and RNA sequencing was performed on Illumina’s NovaSeq platform using a 150 bp paired-end protocol with a depth of >20 million reads per sample.

#### RNA-seq Data Analysis

Quality control (QC) checks on the RNA-seq raw reads were performed using the FastQC tool, followed by read trimming to remove bases with low median (or bottom-quartile) scores. STAR Aligner tool mapped processed RNA-seq reads to the hg38 human genome build. Following data normalization, we applied limma, an empirical Bayesian method, to perform gene expression analysis between groups and detect the differentially expressed (DE) genes. Genes with mean fold-change>|1| and p<0.05 were considered as the threshold for reporting significant differential gene expression. Heatmaps were generated using unsupervised hierarchical clustering (HC) analysis based on the differentially expressed genes (DEGs).

### Single-cell RNA sequencing (scRNA-seq)

Automated single-cell capture and cDNA synthesis were performed on untreated and TAK-715-treated (pre vs. post-treatment) PCa cell lines on the 10X Genomics Chromium platform using the Chromium Next GEM Single Cell 5’ Gene Expression (Dual Index) kit. Single-cell RNA sequencing was performed on the Illumina HiSeq 2500 NGS platform (Paired-end, 2*125bp, 100 cycles, v3 chemistry) at >5 million reads per sample.

### scRNA-seq data analysis

Single-cell RNA-seq datasets were generated as matrices in the Hierarchical Data Format (HDF5 or H5). We used CellRanger, Seurat, and Partek Flow software packages to pre-process and analyze the scRNA-seq data. Highly variable genes were selected for clustering analysis based on a graph-based clustering approach. The visualization of cell populations was performed using T-distributed stochastic neighbor embedding (t-SNE) and Uniform Manifold Approximation and Projection (UMAP) for biomarker-based identification to delineate subclones representing TX-resistant cells, potential TAK-715 target subclones, and cancer stem cell signatures, as well as TAK-715 treatment-induced erosion of these subclones.

### Pathway analysis

The potential biological functions of DEGs were investigated using the Gene Set Enrichment Analysis (GSEA) method to identify biological pathways that share common biological function, chromosomal location, or regulation. The method derives its power by focusing on gene sets, that is, groups of genes that share common biological function, chromosomal location, or regulation^61^. We demonstrate how GSEA yields insights into several cancer-related data sets, including leukemia and lung cancer. Notably, where single-gene analysis finds little similarity between two independent studies of patient survival in lung cancer, GSEA reveals many biological pathways in common. The GSEA method is embodied in a freely available software package, together with an initial database of 1,325 biologically defined gene sets.

Ingenuity pathway analysis (IPA; Qiagen) was performed using top DEGs to reveal the most significantly affected molecular pathways/mechanisms, upstream regulator molecules, downstream effects, biological processes, and causal networks predicted to be activated or inhibited in response to TAK-715 single-agent and combination treatments^62^.

### Quantitative reverse-transcriptase polymerase chain reaction (qRT-PCR)

Cell lines were plated and treated with TAK-715 single-agent, in combination with DTX, or with vehicle (0.5% DMSO) for 24 hours. Total RNA isolation was performed using the RNeasy Plus mini kit. RNA was quantified in Nanodrop 2000 (Thermo Fisher Scientific, Waltham, MA, USA), and cDNA was prepared using QuantiTect Reverse Transcription kit (Qiagen). Following reverse transcription, the transcribed cDNA was added to TaqMan Fast Advanced Master Mix, and quantitative RT-PCR was performed using TaqMan Gene Expression Assays in the CFX96 Touch Real-Time PCR Detection System(Bio-Rad, Hercules, CA).

### Western Blotting

Cells were treated with TAK-715, DTX, and TAK-715+DTX for 48 hours. Following incubation, untreated and treated cells were harvested, washed, trypsinized, and suspended in RIPA lysis & extraction buffer containing 1X protease inhibitor cocktail and 1X phosphatase inhibitor. Protein concentrations were determined using the Bradford reagent with the Quick Start Bovine Serum Albumin Standard, according to the manufacturer’s protocol. Equal amounts of total protein extract were separated by SDS-PAGE (4-20% gel) and transferred onto polyvinylidene fluoride membranes using the OWL HEP-1 semi-dry transfer apparatus (Thermo Fisher Scientific, Inc., Waltham, MA, USA). The membranes were blocked in Tris-buffered saline with 0.1% Tween-20 containing 3% BSA. Subsequently, the cells were incubated with the primary antibodies at 4°C overnight. Immunoreactivity was detected by using a biomarker-specific horseradish peroxidase-conjugated secondary antibody. The protein bands were developed using ECL, and the exposed image was captured using a ChemiDoc™ MP Imaging System (Bio-Rad). Densitometry analysis was performed using ImageJ software. Expression levels of the proteins were normalized to β-actin, which served as an endogenous control.

### Patient datasets

#### PRAD-TCGA data

Clinical, survival, and gene expression data for Patients with Prostate adenocarcinoma (PRAD) were extracted from the Cancer Genome Atlas (TCGA) Data Portal on the Genomic Data Commons (GDC) server (cancergenome.nih.gov). The interactive web-portal Gene Expression Profiling Interactive Analysis (GEPIA) was used for in-depth analysis of TCGA gene expression data files and to compare transcriptome data on target candidate pathway genes with tumor metastasis and patient survival from the prostate expression data matrix^63,64^.

#### Localized prostate cancer dataset

Disease-free survival, biochemical recurrence, and gene expression data on 106 formalin-fixed paraffin-embedded (FFPE) prostatectomy samples on 100 PCa patients from 3 independent sites (the Atlanta VA Medical Center, Moffitt Cancer Center, and Sunnybrook Health Sciences Center at the University of Toronto)) were downloaded from the Gene Expression Omnibus database (Accession number: GSE54460; BioProject ID PRJNA237581)^65^. Analysis of the association of gene expression with DFS was performed using patient RNA-seq data and corresponding clinical information (DFS months and biochemical recurrence). Patients were stratified into high-expression and low-expression groups based on an optimal gene expression cutoff, and Kaplan-Meier survival analysis was performed using the survminer package. Survival differences between groups were assessed using the log-rank test. For each gene, a univariate Cox proportional hazards regression model was fitted using gene expression as a continuous variable and DFS as the outcome. Hazard ratios (HRs), 95% confidence intervals (CIs), and Wald test p-values were calculated using the survival package in R. Genes with p<0.05 and effect size (|log2(HR)| > 1) were considered significantly associated with DFS.

Further, a novel ‘reverse-match’ approach was applied to identify TAK-715-induced gene expression signatures that have the potential to ‘reverse’ the input signature (genes associated with poor survival in patient datasets)^23^.

### Statistical analysis

All statistical analyses were performed using R (the R Project for Statistical Computing and Graphics) and GraphPad Prism. All tests were two-sided, and p<0.05 was considered statistically significant, unless otherwise specified. Multiple-testing correction was performed using the Benjamini-Hochberg method.

## AUTHOR CONTRIBUTIONS

Conceptualization: SC, UKM, and AKM.

Methodology, Investigation & Validation: SC, SM, FH, FA, SB, and TMG.

Data Analysis: SC, PM, UKM, TMG, and AKM;

Visualization: SC, SM, PM, and AKM

Writing: original draft: SC, PM, TMG, and AKM;

Writing: review & editing: RDA, TMG, and AKM;

Supervision & Project administration: AKM.

All authors have read and agreed to the published version of the manuscript.

## ACKNOWLEDGEMENTS

SC and SB were supported by Presidential Fellowship from Auburn University. SC was a graduate student at Auburn University when this work was completed. The current affiliation of SC is Bristol Myers Squibb, Boston, MN. SM was a postdoctoral researcher at Auburn University when this work was completed. The current affiliation of SM is Champions Oncology, Inc., Rockville, MD.

