## Supplementary material for "Dual inhibition of p38α MAPK and Casein kinase δ/ε as a novel treatment strategy for AR-independent and taxane-resistant advanced prostate cancer": Supplementary Figures TAK715_bioRxiv.pdf

**Figure S1. TAK-715 shows single-agent cytotoxicity in androgen-independent mCRPC cell lines.** *In vitro* cell viability profile of Human CRPC cell lines. i) Androgen-independent mCRPC; ii) AR-driven CRPC.

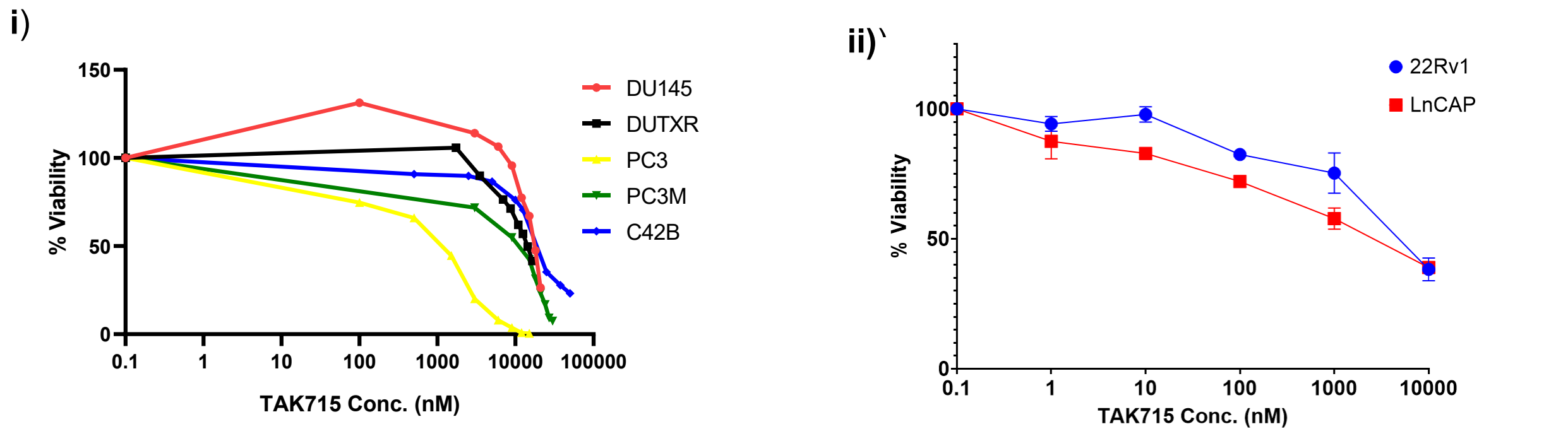

|  | Cell Line | TAK-715 (nM) |
| --- | --- | --- |
| AR-driven CRPC | 22Rv1 | 4813.92 |
|  | LNCaP | 2591.96 |
| AR-independent mCRPC | DU145 | 17624.25 |
|  | DUTXR | 14261.31 |
|  | PC3 | 1245.24 |
|  | PC3M | 11052.63 |
|  | C42B | 19794.35 |
| Mouse | RM1 | 7500.99 |

**Figure S2A. Assessment of cellular morphology following Docetaxel + TAK-715 combination treatment mCRPC cells.** *In vitro* live cell imaging to study the cellular morphology of Control (0.5% DMSO), TAK-715 single agent and DTX+TAK-715 combination treated AR-ve mCRPC cells I-III. Representative figures show the effect of the primary (DTX) and secondary (TAK-715) drugs on cell count and cell morphology of mCRPC cells. The images were captured on the PC3-Luc cells before (0h) and after (48h) TAK-715 treatment either as a single agent or in combination. Microscopy results show significantly higher cell death in combination treatment compared to single-drug treatment for all cell lines; IV. ImageJ data analysis showed a significant difference in cell density for both TAK-715 single-agent and DTX+TAK-715 combination treatments. Results show significantly higher cell death in combination treatment at combination dose compared to single-drug treatment for PC3 cell lines (Significant value \* = $p \leq 0.05$).

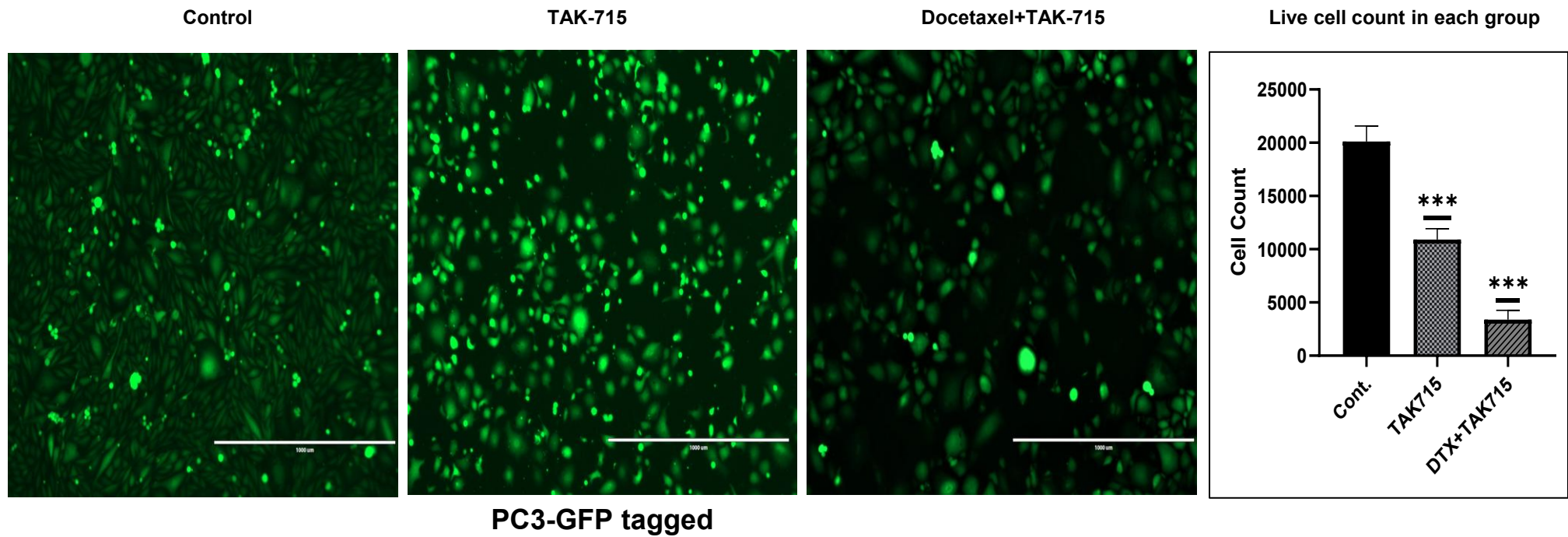

**Figure S2B. Assessment of nuclear morphology.** Representative figures showing the TAK-715-based treatment (single-agent and combination with DTX) on cell nucleus morphology of mCRPC cells. NucBlue Live reagent is frequently used to distinguish condensed nuclei in apoptotic cells. The microscopy images were captured on the PC3-Luc cells before (0h) and after (48h) treatment. Microscope images showing treatment effect on the cell lines PC3. Similar results were obtained for all mCRPC lines.

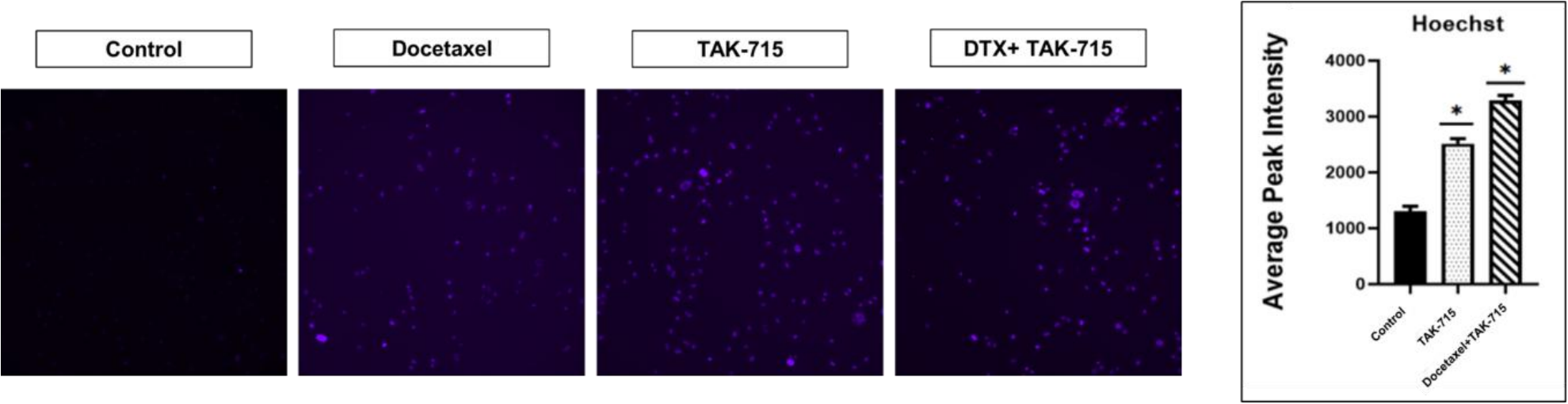

**Figure S3A. Docetaxel + TAK-715 combination enhances apoptosis in mCRPC.** Quantitative measurement of the % apoptotic mCRPC cells (Annexin-V positively stained) exposed to TAK-715 single agent and Docetaxel+TAK-715 combination treatment, using Annexin V/PI staining.

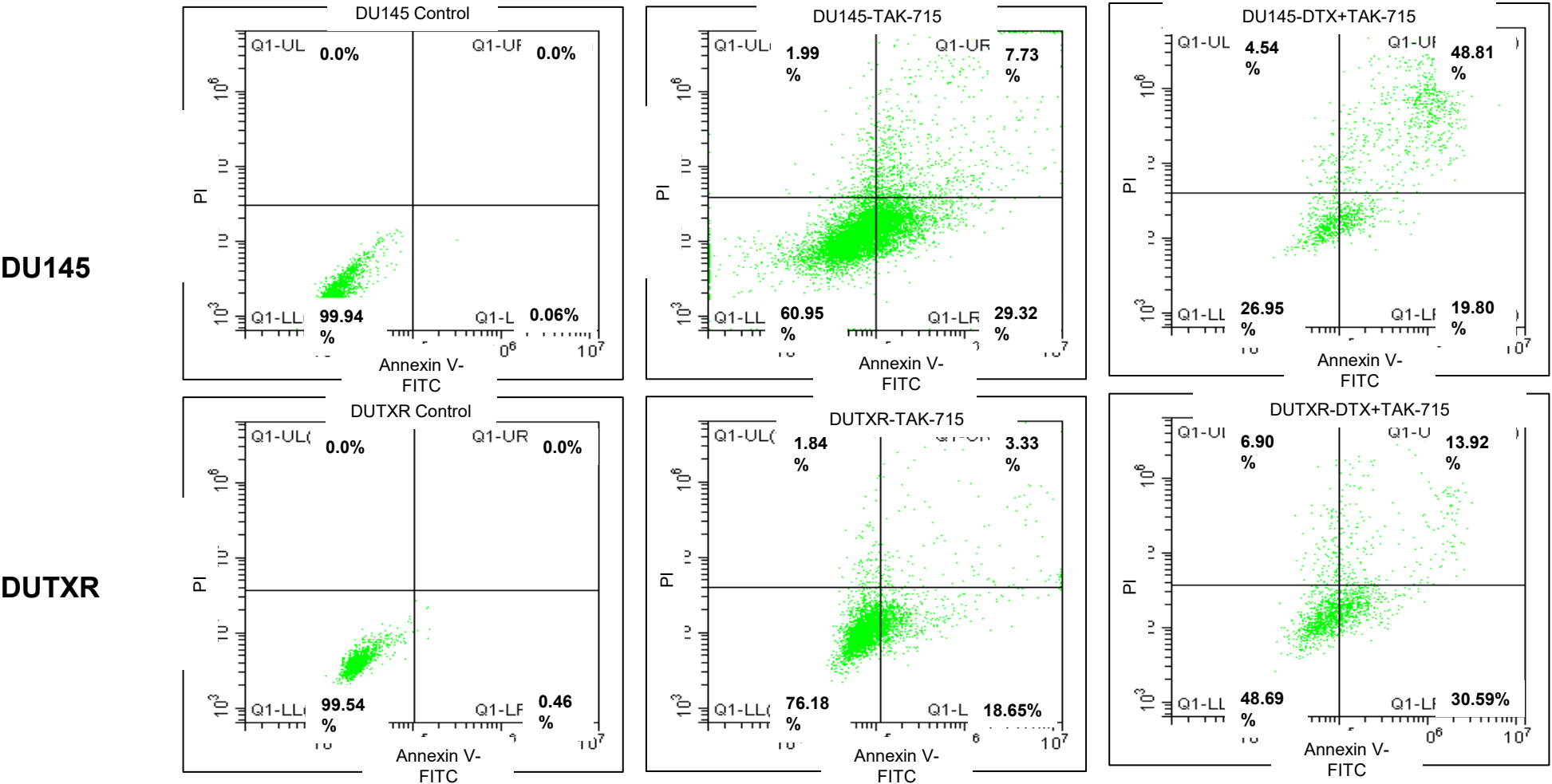

**Figure S3B. Docetaxel + TAK-715 combination enhances apoptosis in mCRPC.** Caspase-3/7 activity assay of Docetaxel and TAK-715 single agent and combination-treated AR<sup>null</sup> mCRPC cell lines. (i) DU145, (ii) DUTXR, (iii) PC-3, (iv) PC-3M. The data shows a higher level of induction of apoptotic pathway in combination treatment compared to single-agent treatment (Significance P-value \* = $p \leq 0.05$). This was further confirmed by Annexin V-FITX/PI-based flow-cytometric apoptosis analysis.

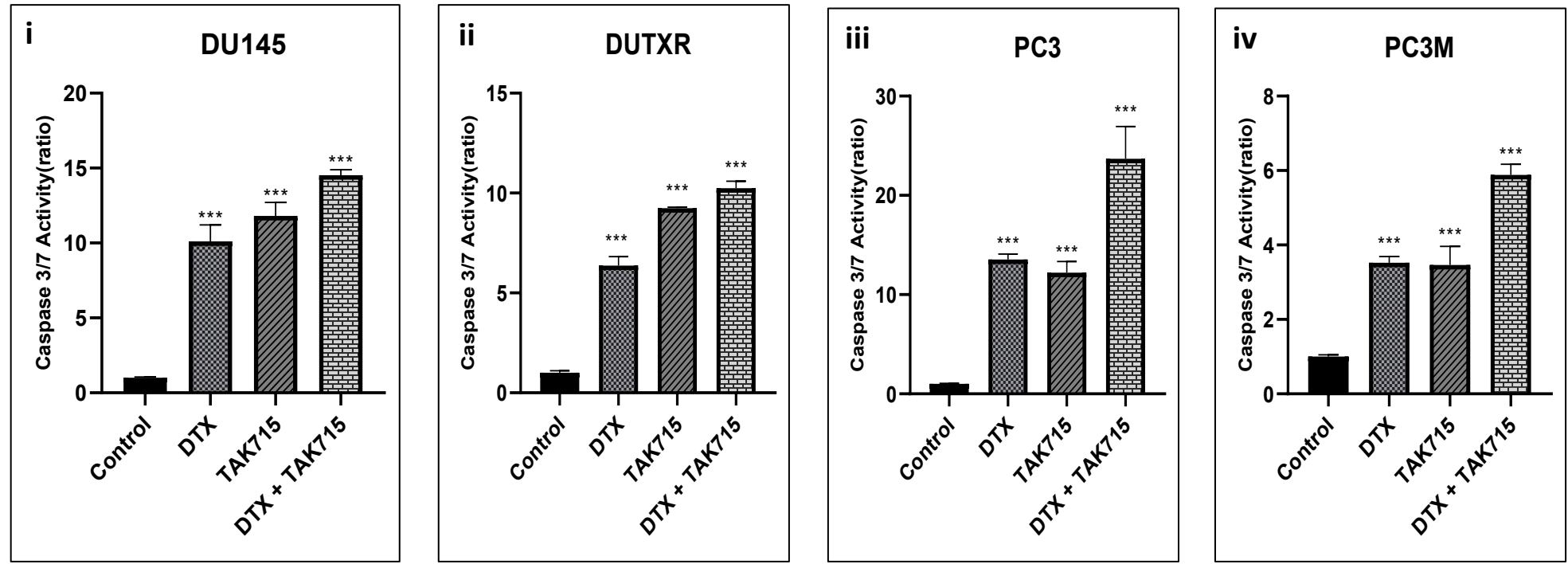

**Figure S4. Gene Set Enrichment Analysis (GSEA)** confirms inflammatory and interferon-responsive transcriptional programs, such as TNFα Signaling via NFκB and IFN-$\alpha/\beta$ (Type I interferon), as the top downregulated pathways enriched in response to TAK-715 treatment.

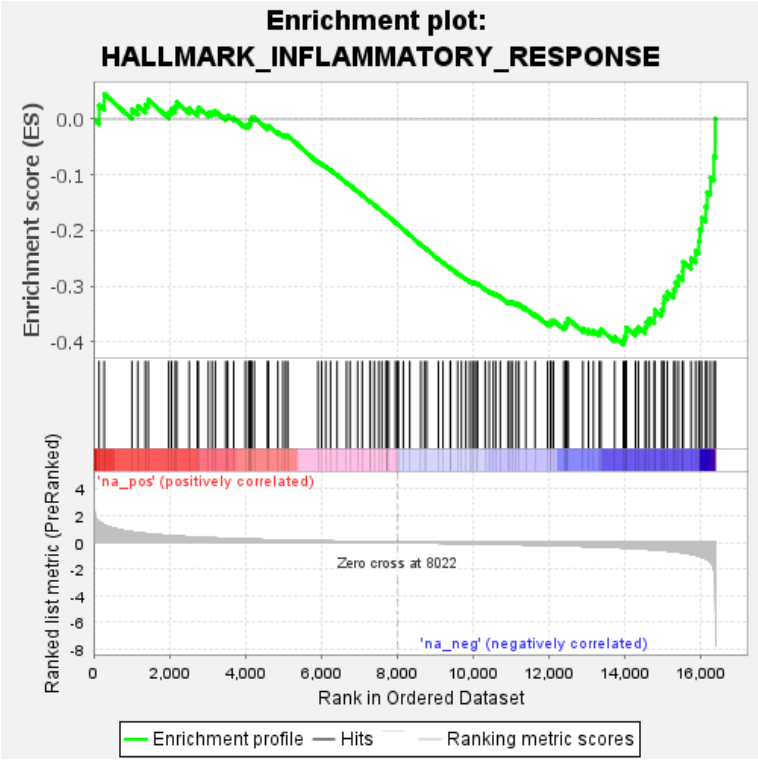

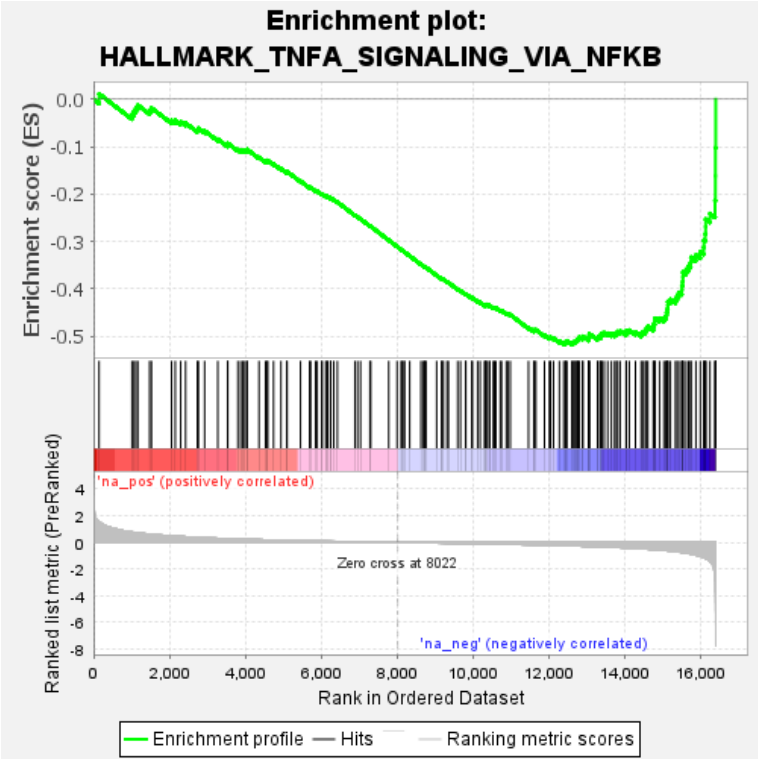

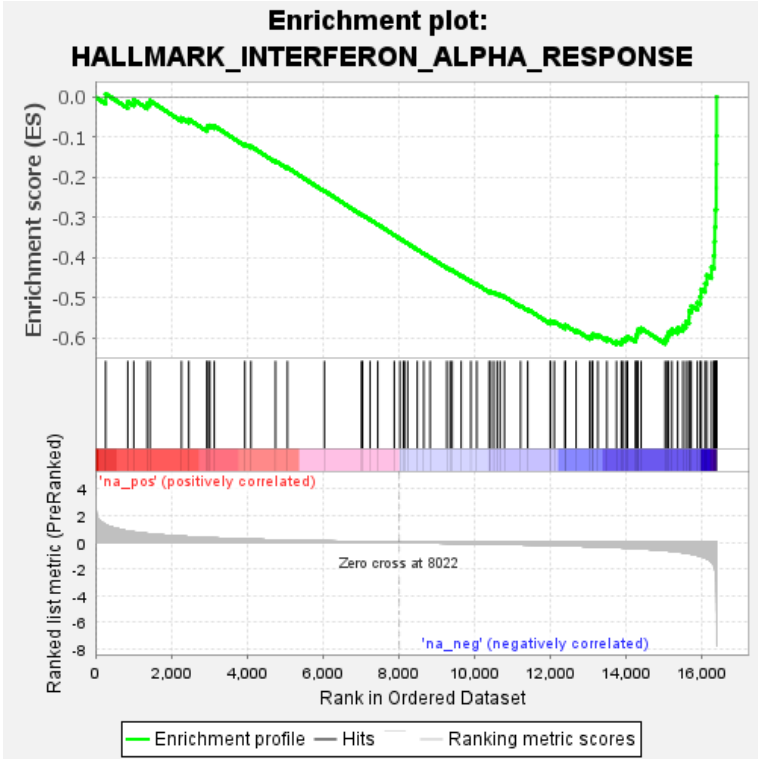

**Figure S5. Combination of Taxane and TAK-715 significantly down-regulated HES1- a transcriptional repressor associated with cancer stemness and multi-drug resistance. TX+TAK-715 combination significantly down-regulates HES1 expression in mCRPC cells. A.** Quantitative qPCR measurement of HES-1 expression in Control vs combination treated AR-ve mCRPC cell lines. **B.** Immunoblotting analysis of HES-1 expression in Control, single agent DTX, single agent TAK-715, and DTX+TAK-715 combination-treated AR-ve mCRPC cell lines.

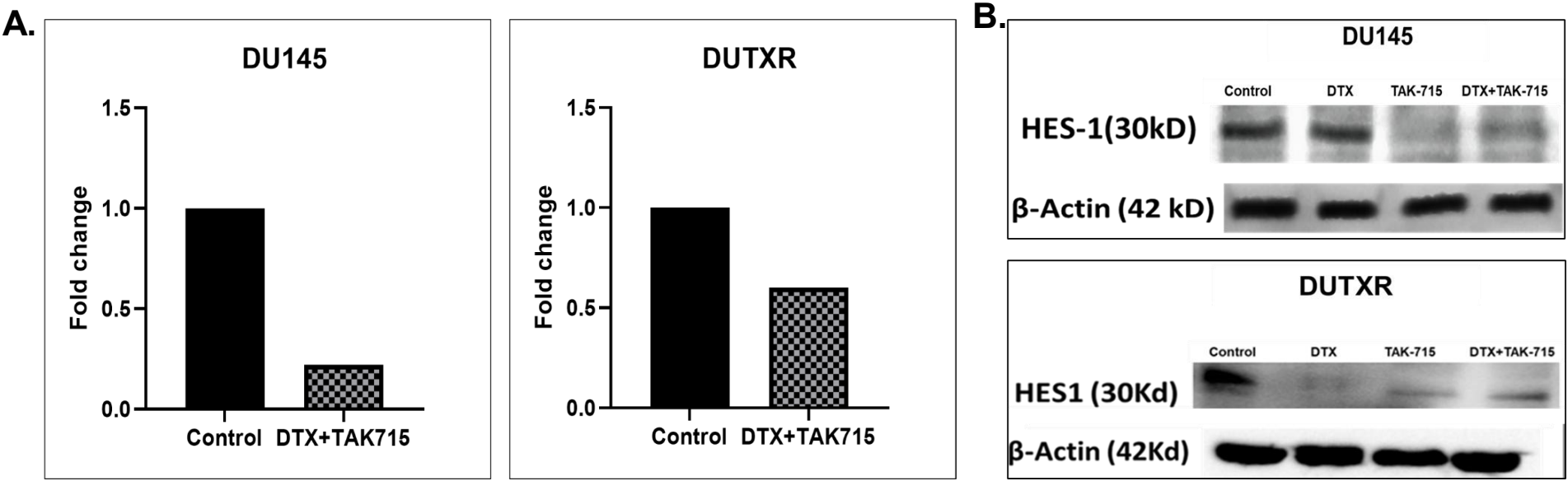

**Figure S6. Comparison of causal networks associated with each treatment group and its validation using *in silico* database analysis. A.** Causal network analysis based on the expression of the genes predicted the up-regulation of miR-132 and the down-regulation of the RICTOR pathway as the top upstream regulator. **B.** GEO prostate cancer dataset (GSE21032) has shown that prostate cancer tissue has a low abundance of miR-132 compared to normal prostate tissue.

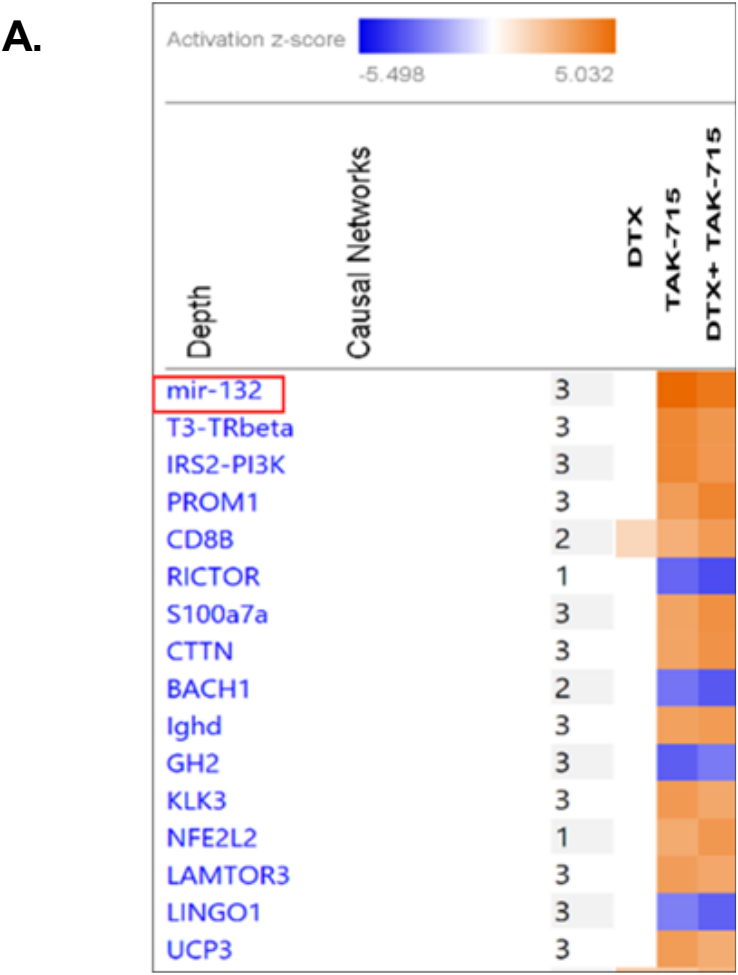

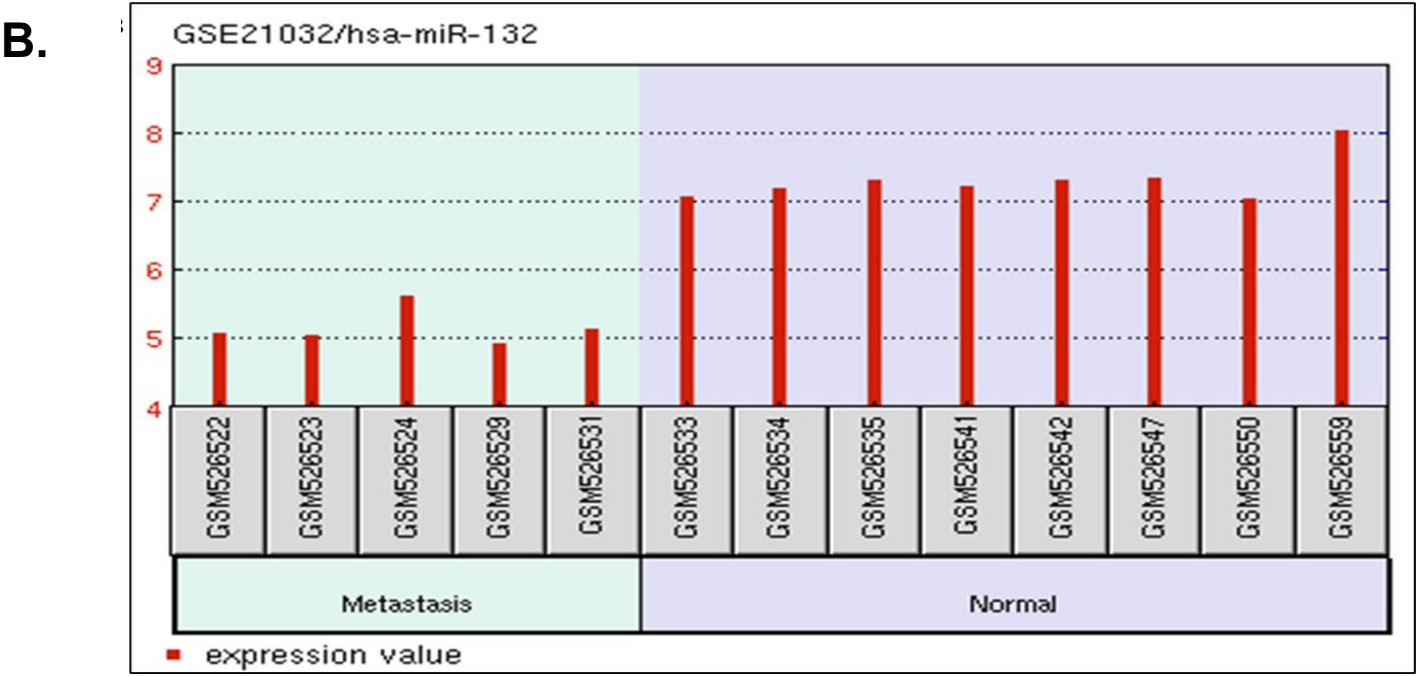

**Figure S7. Single-agent cytotoxicity of Taxanes in mCRPC cell line pair.** *In vitro* cell viability profile of a representative parental TX-sensitive and clonally-derived TX-resistant mCRPC cell line pair. i) Docetaxel (DTX); ii) Cabazitaxel (CBZ).

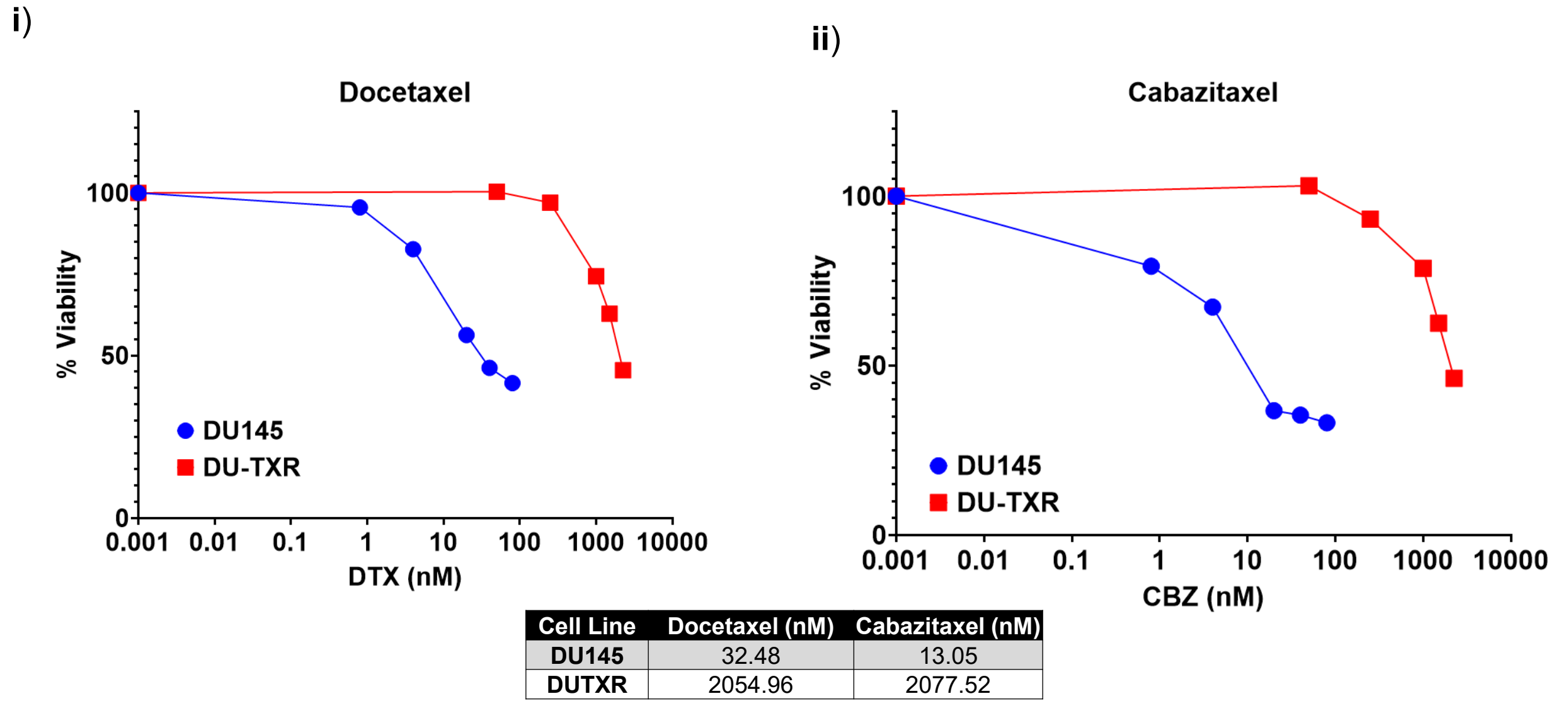
