## Supplementary material for "Dual inhibition of p38α MAPK and Casein kinase δ/ε as a novel treatment strategy for AR-independent and taxane-resistant advanced prostate cancer": Supplementary Tables_TAK715_092626_bioRxiv.pdf

**Supplementary Table S1.** List of cell lines, reagents, and supplies. (ATCC, American Type Culture Collection; BSA, bovine serum albumin; DCFDA, 2',7'-dichlorofluorescin diacetate; ECL, enhanced chemiluminescence; PVDF, polyvinylidene fluoride.)

| Sl. No | Reagent/Material | Manufacturer | Location |
| --- | --- | --- | --- |
| 1 | PC3, DU145, C4-2B, LnCaP, 22Rv1 cell lines | ATCC | Manassas, VA, USA |
| 2 | DUTXR taxane-resistant prostate cancer cell line | Provided by Dr. Feng Li | Auburn University, Auburn, AL, USA |
| 3 | MTT (3-(4,5-dimethylthiazol-2-yl)-2,5-diphenyltetrazolium bromide) | Sigma-Aldrich | St. Louis, MO, USA |
| 4 | DCFDA | Sigma-Aldrich | St. Louis, MO, USA |
| 5 | JC-1 | Sigma-Aldrich | St. Louis, MO, USA |
| 6 | Docetaxel | Selleck Chemicals | Houston, TX, USA |
| 7 | Cabazitaxel | Selleck Chemicals | Houston, TX, USA |
| 8 | TAK-715 | Selleck Chemicals | Houston, TX, USA |
| 9 | Caspase-Glo® 3/7 Assay System | Promega Corporation | Madison, WI, USA |
| 10 | FITC Annexin V Apoptosis Detection Kit I | BD Biosciences | San Jose, CA, USA |
| 11 | FxCycle™ PI/RNase Staining Solution | Thermo Fisher Scientific | Waltham, MA, USA |
| 12 | Comet Assay Kit | Trevigen, Inc. | Gaithersburg, MD, USA |
| 13 | RNeasy Plus Mini Kit | QIAGEN | Hilden, Germany |
| 14 | QuantiTect Reverse Transcription Kit | QIAGEN | Hilden, Germany |
| 15 | TaqMan Fast Advanced Master Mix | Thermo Fisher Scientific | Foster City, CA, USA |
| 16 | TaqMan Gene Expression Assays | Thermo Fisher Scientific | Foster City, CA, USA |
| 17 | RIPA Lysis Buffer | Thermo Fisher Scientific | Waltham, MA, USA |
| 18 | Protease Inhibitor Cocktail | Thermo Fisher Scientific | Waltham, MA, USA |
| 19 | Phosphatase Inhibitor | Thermo Fisher Scientific | Waltham, MA, USA |
| 20 | Bradford Reagent | Thermo Fisher Scientific | Waltham, MA, USA |
| 21 | Pierce™ ECL Western Blotting Substrate | Thermo Fisher Scientific | Waltham, MA, USA |
| 22 | Quick Start Bovine Serum Albumin Standard | Bio-Rad Laboratories | Hercules, CA, USA |
| 23 | Tris Buffer | Bio-Rad Laboratories | Hercules, CA, USA |
| 24 | 10% Tween 20 Solution | Bio-Rad Laboratories | Hercules, CA, USA |
| 25 | Polyvinylidene Fluoride (PVDF) Membrane | EMD Millipore | Billerica, MA, USA |
| 26 | Bovine Serum Albumin (BSA) | VWR International | Radnor, PA, USA |
| 27 | 6-well Cell Culture Plates | VWR International | Radnor, PA, USA |
| 28 | 24-well Cell Culture Plates | VWR International | Radnor, PA, USA |
| 29 | 96-well Cell Culture Plates | VWR International | Radnor, PA, USA |
| 30 | Primary Antibodies | Cell Signaling Technology | Danvers, MA, USA |
| 31 | Secondary Antibodies | Cell Signaling Technology | Danvers, MA, USA |
